# Swarm Learning as a privacy-preserving machine learning approach for disease classification

**DOI:** 10.1101/2020.06.25.171009

**Authors:** Stefanie Warnat-Herresthal, Hartmut Schultze, Krishnaprasad Lingadahalli Shastry, Sathyanarayanan Manamohan, Saikat Mukherjee, Vishesh Garg, Ravi Sarveswara, Kristian Händler, Peter Pickkers, N. Ahmad Aziz, Sofia Ktena, Christian Siever, Michael Kraut, Milind Desai, Bruno Monnet, Maria Saridaki, Charles Martin Siegel, Anna Drews, Melanie Nuesch-Germano, Heidi Theis, Mihai G. Netea, Fabian Theis, Anna C. Aschenbrenner, Thomas Ulas, Monique M.B. Breteler, Evangelos J. Giamarellos-Bourboulis, Matthijs Kox, Matthias Becker, Sorin Cheran, Michael S. Woodacre, Eng Lim Goh, Joachim L. Schultze, German COVID-19 OMICS Initiative (DeCOI)

## Abstract

Identification of patients with life-threatening diseases including leukemias or infections such as tuberculosis and COVID-19 is an important goal of precision medicine. We recently illustrated that leukemia patients are identified by machine learning (ML) based on their blood transcriptomes. However, there is an increasing divide between what is technically possible and what is allowed because of privacy legislation. To facilitate integration of any omics data from any data owner world-wide without violating privacy laws, we here introduce Swarm Learning (SL), a decentralized machine learning approach uniting edge computing, blockchain-based peer-to-peer networking and coordination as well as privacy protection without the need for a central coordinator thereby going beyond federated learning. Using more than 14,000 blood transcriptomes derived from over 100 individual studies with non-uniform distribution of cases and controls and significant study biases, we illustrate the feasibility of SL to develop disease classifiers based on distributed data for COVID-19, tuberculosis or leukemias that outperform those developed at individual sites. Still, SL completely protects local privacy regulations by design. We propose this approach to noticeably accelerate the introduction of precision medicine.

## Introduction

Fast and reliable detection of patients with severe illnesses is a major goal of precision medicine^1^. The measurement of molecular phenotypes for example by omics technologies^2^ and the application of sophisticated bioinformatics including artificial intelligence (AI) approaches^3–7^ opens up the possibility for physicians to utilize large-scale data for diagnostic purposes in an unprecedented way. Yet, there is an increasing divide between what is technically possible and what is allowed because of privacy legislation^8^ (hhs.gov, https://www.hhs.gov/hipaa/index.html, 2020; Intersoft Consulting, General Data Protection Regulation, https://gdpr-info.eu; Convention for the Protection of Individuals with regard to Automatic Processing of Personal Data, https://rm.coe.int/16808ade9d). Particularly, in a global health crisis, as in the case of the infection with severe acute respiratory syndrome coronavirus 2 (SARS-CoV-2) leading to the pandemic spread of coronavirus disease 2019 (COVID-19)^9–11^, reliable, fast, secure and privacy-preserving technical solutions based on AI principles are now believed to add to the armamentarium to quickly answer important questions in the fight against such threats^12–15^. These AI-based concepts range from protein structure prediction^16^, drug target prediction^17^, knowledge sharing^18^, tools for population control^19, 20^ to the assistance of healthcare personnel, e.g. by developing AI-based coronavirus diagnostic software^21, 22^. Considering the more clinically oriented AI-based technical solutions, any such progress might also induce improvements for a variety of deadly diseases including other major infections or cancer^23^. For example, the principles of a recently introduced AI-system for diagnosing COVID-19 pneumonia and predicting disease outcome using computed tomography^22^ might be further developed to identify patients with tuberculosis or lung cancer in the future^24^. At the same time, we need to consider important standards relating to data privacy and protection, such as Convention 108(+) of the Council of Europe (Convention for the Protection of Individuals with regard to Automatic Processing of Personal Data, https://rm.coe.int/16808ade9d), which regulate the use and sharing of health data including in AI-based approaches, irrespective of the occurrence of a pandemic crisis.

AI-based solutions intrinsically rely on appropriate algorithms^25^, but even more so on large enough datasets for training purposes^26^. Since the domain of medicine is inherently decentralized, the volume of data available locally is often insufficient to train reliable classifiers^27–29^. As a consequence, centralization of data, for example via cloud solutions, has been one model to address the local limitations^30–32^. While beneficial from an AI-perspective, centralized solutions were shown to have other inherent hurdles, including increased data traffic of large medical data, data ownership, privacy and security concerns when ownership is disconnected from access and usage curation and thereby creating data monopolies favoring data aggregators^26^. Consequently, solutions to the challenges of central data models in AI - particular when dealing with medical data - must be effective, with high accuracy and efficiency, privacy- and ethics-preserving, secure, and fault-tolerant by design^33–36^. Federated AI has been introduced to address some of these aspects^26, 37–39^. While data are kept locally (at the edge) and privacy issues are addressed^40, 41^, the model parameters in federated AI are still handled by central custodians who as the intermediaries concentrate power of the learning to themselves. Furthermore, such star-shaped architectures decrease fault tolerance.

We hypothesized that completely decentralized AI solutions overcome current technical shortcomings and at the same time accommodate for inherently decentralized data structures in medicine as well as pronounced data privacy and security regulations. The solution would 1) need to keep large medical data locally with the data owner, 2) require no raw data exchange thereby also reducing data traffic and issues related to central storage, 3) provide high level data security and privacy protection, 4) guarantee secure, transparent and fair onboarding of decentralized members participating in the learning network without the need for a central custodian, 5) allow for parameter merging with equal rights for all members requiring no central custodian, and 6) protect the ML models from attacks. To address these points, we introduce the concept of Swarm Learning (SL). SL combines decentralized hardware infrastructures, distributed ML technique based on standardized AI engines with a permissioned blockchain to securely onboard members, dynamically elect the leader among the members, and merge model parameters. All processes are orchestrated by an SL library and an iterative learning procedure applying AI solutions to compute problems with decentralized private data.

Medicine is a prime example to illustrate the advantages of this AI approach. Without any doubt, numerous medical features including radiograms or computed tomographies, proteomes, metagenomes or microbiomes derived from body fluids including nasal or throat swaps, blood, urine or stool are all excellently suitable medical data for the development of AI-based diagnostic or outcome prediction classifiers. We here chose to evaluate the cellular compartment of peripheral blood, either in form of peripheral blood mononuclear cells (PBMC) or whole blood-derived transcriptomes, since blood-derived transcriptomes include important information about the patients’ immune response during a certain disease, which in itself is an important molecular information^42, 43^. In other words, in addition to the use of blood-derived high-dimensional molecular features for a diagnostic or outcome classification problem, blood transcriptomes could be further utilized in the clinic to systematically characterize ongoing pathophysiology, predict patient-specific drug targets and trigger additional studies targeting defined cell types or molecular pathways, making this feature space even more attractive to answer a wide variety of medical questions. Here, we illustrate that newly generated blood transcriptome data together with data derived from more than 14,000 samples in more than 100 studies combined with AI-based algorithms in a Swarm Learning environment can be successfully applied in real-world scenarios to detect patients with leukemias, tuberculosis or active COVID-19 disease in an outbreak scenario across distributed datasets without the necessity to negotiate and contractualize data sharing.

## Results

### Concept of Swarm Learning

Machine learning (ML) of any data including genome or transcriptome data requires the availability of sufficiently large datasets^26, 44^ and the respective compute infrastructure including data storage for data processing and analytics^45^. Conceptually, if data and compute infrastructure is sufficiently available locally, ML can be performed locally (‘at the edge’) (**Fig. 1a**). However, often medical data are not sufficiently large enough locally and similar approaches are performed at different locations in a disconnected fashion. These limitations have been overcome by cloud computing where data are moved centrally to perform training of ML algorithms in a centralized compute environment (**Fig. 1b**). Compared to local approaches, cloud computing can significantly increase the amount of data for training ML algorithms and therefore significantly improve their results^26^. However, cloud computing has other disadvantages such as data duplication from local to central data storage, increased data traffic and issues with locally differing data privacy and security regulations^46^. As an alternative, federated cloud computing approaches such as Google’s federated learning^38^ and Facebook’s elastic averaging SGD (Deep learning with Elastic Averaging SGD, http://papers.neurips.cc/paper/5761-deep-learning-with-elastic-averaging-sgd.pdf) have been developed. In these models, dedicated parameter servers are responsible for aggregating and distributing local learning (**Fig. 1c**). A disadvantage of such star-shaped system architectures is the remainder of a central structure, which hampers implementation across different jurisdictions and therefore still requires the respective legal negotiations. Furthermore, the risk for a single point of failure at the central structure reduces fault-tolerance.

**Figure 1.**
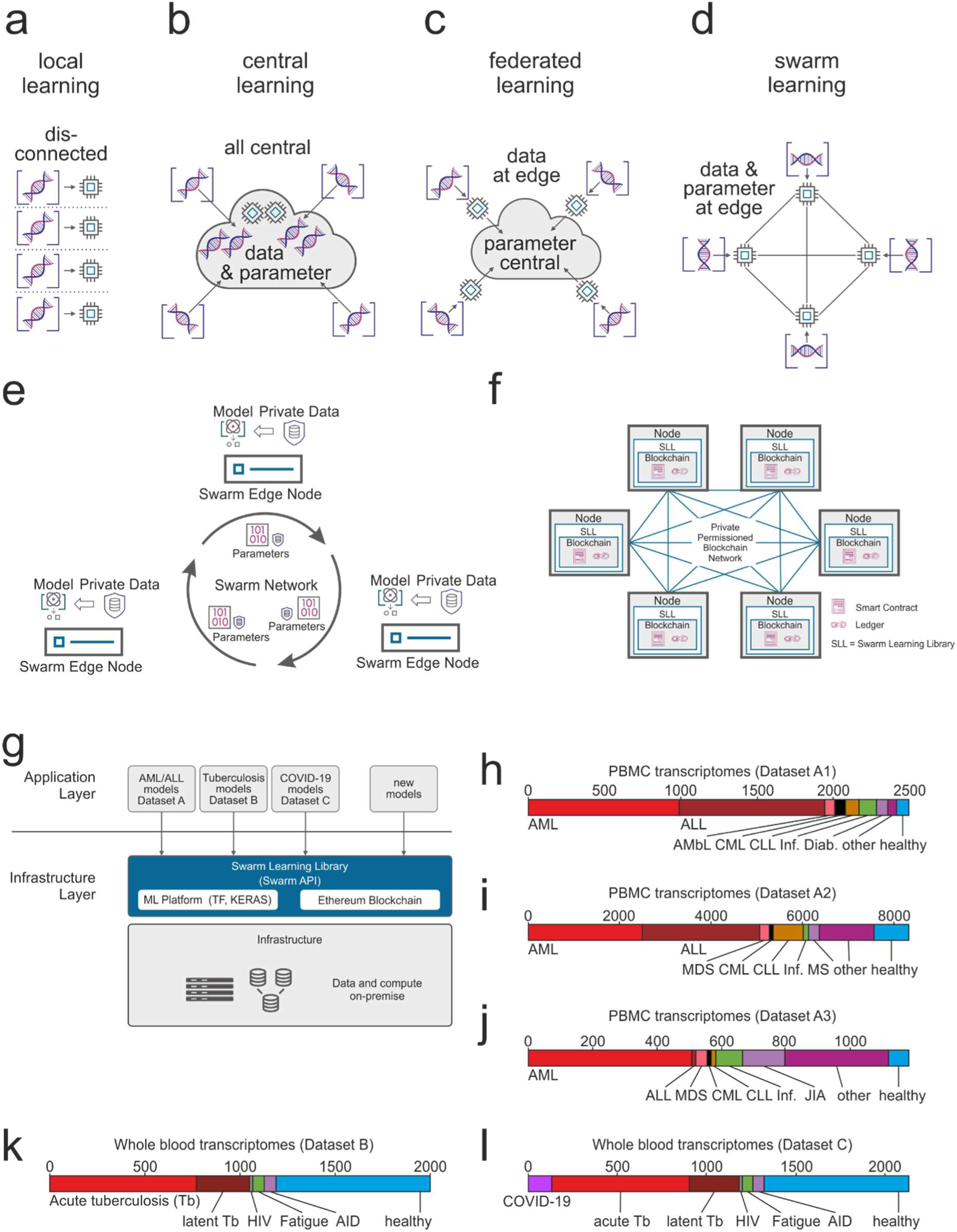
Concept of Swarm Learning. **(a-d)** The principles of Swarm Learning in contrast to other machine learning concepts. **(a)** Illustration of the concept of local learning with data and computation at different, but disconnected locations. **(b)** Principle of cloud-based machine learning where data from contributing centers move copies of the data to a central cloud-based storage; centrally located data are then used for central - often cloud-based - machine learning. **(c)** Federated learning with data being kept with the data contributor and computing is also performed at the site of local data storage and availability, yet parameter settings of machine learning are orchestrated by a central parameter server. **(d)** Swarm Learning principle with swarm nodes being connected in a democratic fashion (enabled by blockchain technology) without the need for a central custodian or parameter server. Data privacy is preserved, data is kept where it is generated, computation is achieved locally and learning parameters are shared within the Swarm Network. **(e)** Schematic representation of the Swarm Network consisting of the Swarm Edge Nodes (short ‘nodes’) that exchange parameters for learning, which is implemented using blockchain technology. Use of private data at each node together with the model provided via Swarm Network. **(f)** Concept and outline of the private permissioned blockchain network as a layer of the Swarm Learning network. Each node consists of the blockchain, including the ledger and smart contract, as well as the Swarm Learning Library (SLL) with the API to interact with other nodes within the network. **(g)** Application and infrastructure layer as part of the Swarm Learning concept. **(h-l)** Description of the transcriptome datasets used within this study: Dataset **(h)** A1 and **(i)** A2, two microarray-based transcriptome datasets of peripheral blood mononuclear cells (PBMC). **(j)** Dataset A3, RNA-seq based transcriptomes of PBMC. Dataset **(k)** B and **(l)** C, RNA-seq based whole blood transcriptome datasets. Abbreviations: *AML*, Acute Myeloid Leukemia; *ALL*, Acute Lymphoblastic Leukemia; *COVID-19*, CoronaVirus Disease 2019; *API*, Application Programming Interface; ML, Machine Learning; TF, Tensor Flow; KERAS, Open Source Deep Learning Library; *AMbl*, Acute Myeloblastic Leukemia; *CML*, Chronic Myeloid Leukemia; *CLL*, Chronic Lymphocytic Leukemia; *Inf*., Infections, *Diab*., Diabetes Type II; *MDS*, Myelodysplastic Syndrome; *MS*, multiple sclerosis; *JIA*, Juvenile idiopathic arthritis; *Tb*, tuberculosis; *HIV*, Human Immunodeficiency Virus, *AID*, Acute Infectious Disease. SLL Swarm Learning Library.

In an alternative model, which we introduce here as Swarm Learning (SL), we dismiss the dedicated server and allow parameters and models to be shared only locally (**Fig. 1d**). While parameters are shared via the swarm network, the models are built independently on private data at the individual sites, here referred to as swarm edge nodes (short ‘nodes’) (**Fig. 1e**). SL provides security measures to guarantee data sovereignty, security and privacy realized by a private permissioned blockchain technology which enables different organizations or consortia to efficiently collaborate (**Fig. 1f**). In a private permissioned blockchain network, each participant is well defined and only pre-authorized participants can execute the transactions. Hence, they use computationally inexpensive consensus algorithms, which offers better performance and scalability. Onboarding of new members or nodes can be done dynamically with the appropriate authorization measures to know the participants of the network, which allows continuous scaling of learning (**Extended Data Fig. 1a**). A new node enrolls via a blockchain smart contract, obtains the model, and performs local model training until defined conditions for synchronization are met. Next, model parameters are exchanged via a Swarm API with the rest of the swarm members and merged for an updated model with updated parameter settings to start a new round of training at the nodes. This process is repeated until stopping criterions are reached, which are negotiated between the swarm nodes/members. The leader is dynamically elected using a blockchain smart contract for merging the parameters and there is no need for a central coordinator in this swarm network. The parameter merging algorithm is executed using a blockchain smart contract thus protects it from semi-honest or dishonest participants. The parameters can be merged by the leader using different functions including average, weighted average, minimum, maximum, or median functions. The various merge techniques and merge frequency enables SL to efficiently work with imbalanced and biased data. As currently developed, SL works with parametric models with finite sets of parameters, such as linear regression or neural network models.

At each node, SL is conceptually divided into infrastructure and application layer (**Fig. 1g**). On top of the physical infrastructure layer (hardware) the application environment contains the ML platform, the blockchain, and the SL library (SLL) including the Swarm API in a containerized deployment, which allows SL to be executed in heterogeneous hardware infrastructures (**Fig. 1g, Supplementary Information**). The application layer consists of the content, the models from the respective domain, here medicine (**Fig. 1g**), for example blood transcriptome data from patients with leukemias, tuberculosis and COVID-19 (**Fig. 1h-l**). Collectively, Swarm Learning allows for a completely decentralized and therefore democratized, secure, privacy-preserving, hardware-independent, and scalable machine learning environment, applicable to many scenarios and domains, which we demonstrate with three medical examples.

### Swarm learning robustly predicts leukemias from peripheral blood mononuclear cell data

As a first use case, we chose transcriptomes derived from peripheral blood mononuclear cells (PBMC) of more than 12,000 individuals (**Fig. 1h-j**) separated into three individual datasets (A1, A2, A3) based on the technology used for generating the transcriptomes (2 different microarrays, RNA-seq)^47^. We used a deep neural network (Keras, https://keras.io/, 2015) as the machine learning approach in all three use cases. To assess performance metrics of SL, we simulated scenarios by dividing up the individual samples derived from several independently performed studies (see Material and Methods) within each of the three datasets into non-overlapping training and test sets. The training sets were then distributed to three nodes for training and classifiers were tested at a fourth node (independent test set) (**Fig. 2a**). By assigning the training data to the nodes in different distributions, we mimicked several clinically relevant scenarios (**Supplementary Table 1)**. As cases, we first used samples defined as acute myeloid leukemia (AML), all other samples are termed ‘controls’. Each node within this simulation could stand for a large hospital or center, a network of hospitals performing individual studies together, a country or any other independent institutional organization generating such medical data with local privacy requirements.

**Figure 2.**
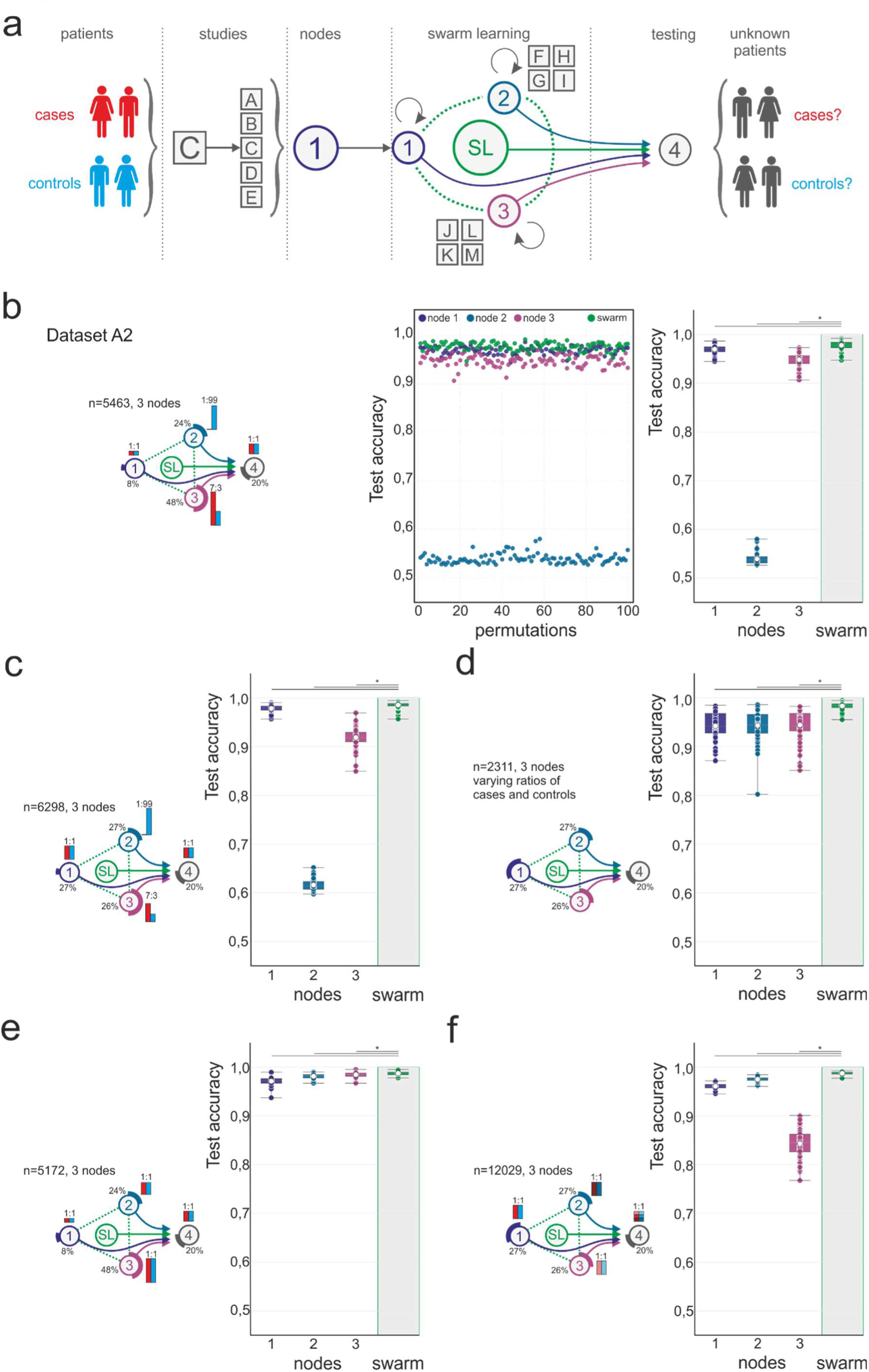
Swarm learning to predict leukemias from PBMC data. **(a)** Schematic representation of the use of the transcriptome data derived from more than 12,000 individuals in over 100 individual studies^47^. Principle of distribution of data to individual Swarm Edge Nodes (short ‘nodes’). Nodes 1-3 were used for training, node 4 for testing. Swarm Learning (SL) was achieved by integrating nodes 1-3 for training following procedures described in detail in Supplementary Information. **(b)** Scenario using Dataset A2. Left panel illustrating the setting of the scenario concerning distribution of cases and controls to individual nodes, as well as total number of samples used for this scenario. Cases (red bar) and controls (blue bar) were distributed unevenly among nodes, the number of samples distributed to each node was also uneven in this scenario. Middle panel shows results of accuracy of all 100 permutations performed for the 3 training nodes individually as well as the results obtained by SL. Accuracy is defined for the independent fourth node used for testing only. Right panel represents box-whisker plot representation of the individual data presented in the middle panel showing mean, 1st and 3rd quartile and whisker type Min/Max. **(c)** Scenario with uneven numbers of cases and controls at the different training nodes but similar numbers of samples at each node to determine impact of these changes on SL performance. Left panel: schematic representation of scenario and right panel: results obtained for accuracy at the test node (node 4) for each of the three training nodes 1-3 and SL independently as box and whisker plot with the same parameter as described for (b). **(d)** Scenario with even numbers at each of the nodes, schematic representation (left panel) and visualization of results as box-whisker plots as in (b) and (c). **(e)** Scenario with even distribution of cases and controls at each training node, but different numbers of samples at each node and overall increase in numbers of samples. Representation of schema and data visualization as in (b-d). **(f)** Scenario where each node obtained samples from different Datasets (node 1: Dataset A1, node 2: Dataset A2, node 3, Dataset A3). Node 4 obtained samples from each Dataset A1-A3 to define impact on technical bias on Swarm Learning performance. Representation of schema and data visualization as in (b-e). Statistical differences between results derived by SL and individual nodes including all permutations performed were calculated with Wilcoxon signed rank test with continuity correction; asterisk and line: p<0.05.

In a first scenario, we randomly distributed samples per node as well as cases and controls unevenly at the nodes and between nodes (dataset A2) (**Fig. 2b**). Sample distribution between sample sets was permuted 100 times (**Fig. 2b, middle panel**) to determine the influence of individual samples on overall performance. Among the nodes, the best test results were obtained by node one with a mean accuracy of 97.0%, mean sensitivity of 97.5% and mean specificity of 96.3% with an even distribution between cases and controls, albeit this node had the smallest number of overall training samples. Node 2 did not produce any meaningful results, which was due to a too low ratio of cases to controls (1:99) for training. Surprisingly, node 3 with the largest number of samples, but an uneven distribution (70% cases : 30% controls) performed worse than node 1 with a mean balanced accuracy of 95.1%. Most importantly, however, SL outperformed each of the nodes resulting in a higher test accuracy in 97.0% of all permutations (mean balanced accuracy 97.7%) (**Fig. 2b, right panel, Supplementary Table 4**). The balanced accuracy of SL was significantly higher (p < 0.001) when compared to the performance of each of the three nodes, despite the fact that information from the poorly performing node 2 was integrated. We also calculated this scenario in datasets A1 and A3 and obtained rather similar results strongly supporting that the performance improvement of SL over single nodes is independent of data collection (studies) and even experimental technologies (microarray (datasets A1, A2), RNA-seq (dataset A3) used for data generation (**Extended Data Fig. 2**).

To test whether more evenly distributed samples at the nodes would improve individual node performance, we distributed similar numbers of samples to each of the nodes but kept case:control ratios as in scenario 1 (**Fig. 2c, Extended Data Fig. 3**). While there was a slight increase in test accuracy at nodes 1 and 2, node 3 performed worse with also higher variance. More importantly, SL still resulted in the best performance metrics (mean 98.5% accuracy) with slightly but significantly (p<0.001) increasing performance compared to the first scenario. Results derived from datasets A1 and A3 echoed these findings (**Extended Data Fig. 3**).

In a third scenario, we distributed the same number of samples across all three nodes, but increased potential batch effects between nodes, by distributing samples of a clinical study independently performed and published in the past only to a dedicated training node. In this scenario, cases and control ratios varied between nodes and left out samples (independent samples) from the same published studies were combined for testing at node 4. Performance of the three nodes was very comparable, but never reached SL results (mean 98.3% accuracy, swarm outperformed all nodes with p<0.001, **Fig. 2d., Extended Data Fig. 4b, Supplementary Data Table 4**), which was also true for datasets A1 and A3 (**Extended Data Fig. 4c-d**). Even when further increasing batch effects by distributing samples from independent published studies to the test node, which means that training and test datasets come from studies performed and published independently, SL outperformed the individual nodes, albeit the variance in the results was increased both at each node and for SL, indicating that study design has an overall impact on classifier performance and that this is still seen in SL (mean 95.6% accuracy, **Extended Data Fig. 4e**).

In a fourth scenario, we further optimized the nodes by increasing the overall sample size at node 3 and keeping case:control ratios even at all nodes (**Fig. 2e, Extended Data Fig. 5a-d**). Clearly, node performance further improved with little variance between permutations, however, even under these ‘node-optimized’ conditions, SL led to higher performance parameters.

In a fifth scenario, we tested whether or not SL was ‘immune’ against the impact of the data generation procedure (microarray versus RNA-seq) (**Fig. 2f, Extended Data Fig. 5e,f**). We recently demonstrated that classifiers trained on data derived by one technology (e.g. microarrays) do not necessarily perform well on another (e.g. RNA-seq)^47^. To test this influence on SL, we distributed the samples from the three different datasets (A1-A3) to one node each, e.g. dataset A1 was used for training only at node 1. We used 20% of the data (independent non-overlapping to the training data) from each dataset (A1-A3) and combined them to form the test set (node 4). Node 3, trained on RNA-seq data, performed poorly on the combined dataset due to the fact that two-thirds of the data in the test set were microarray-derived data. Nodes 1 and 2 performed reasonably well with mean accuracies of 96.1% (node 1) and 97.5% (node 2), however did not reach the test accuracy of SL (98.8%), which also indicated that SL is much more robust toward effects introduced by different data production technologies in transcriptomics (**Fig. 2f, Extended Data Fig. 5e,f**).

Finally, we repeated several of these scenarios with acute lymphoblastic leukemia (ALL) as the second most prevalent disease in dataset A2 (**Extended Data Fig. 6** and data not shown) and demonstrated very similar results with SL outperforming the classifiers built at the nodes. Collectively, these simulations using real-world transcriptome data collected from more than 100 individual studies illustrate that SL would not only allow data to be kept at the place of generation and ownership, but it also outperforms every individual node in numerous scenarios, even in those with nodes included that cannot provide any meaningful classifier results.

### Swarm learning to identify patients with tuberculosis

In infectious diseases, heterogeneity may be more pronounced compared to leukemia, therefore we built a second use case predicting cases with tuberculosis (Tb) from full blood transcriptomes. Of interest, previous work in smaller studies had already suggested that acute tuberculosis or outcome of tuberculosis treatment can be revealed by blood transcriptomics ^48–52^. To apply SL, we generated a new dataset based on full blood transcriptomes derived by PaxGene blood collection followed by bulk RNA-sequencing. We also generated new blood transcriptomes and added existing studies to the dataset compiling a total of 1,999 samples from nine individual studies including 775 acute and 277 latent Tb cases (**Fig. 1k, Extended Data Fig. 7a, Supplementary Table 2**). These data are more challenging, since infectious diseases show more variety due to biological differences with respect to disease severity, phase of the disease or the host response. But also the technology itself is more variable with numerous different approaches for full blood transcriptome sample processing, library production and sequencing, which can introduce technical noise and batches between studies. As a first scenario, we used all Tb samples (latent and acute) as cases and divided Tb cases and controls evenly among the nodes (**Extended Data Fig. S7a-b, Supplementary Table 1**). Similar to AML and ALL, in detecting Tb, SL outperformed the individual nodes in accuracy (mean 93.4%), sensitivity (mean 96.0%) and specificity (mean 90.9%) (**Extended Data Fig. S7b**). To increase the challenge, we decided to assess prediction of acute Tb cases only. In this scenario, latent Tb are not treated as cases but rather as controls (**Extended Data Fig. S7a**). For the first scenario, we kept cases and controls even at all nodes but further reduced the number of training samples (**Fig. 3a-b**). As expected in this more challenging scenario, distinguishing acute Tb from the control cohort (including latent Tb samples), overall performance (mean balanced accuracy 89.1%, mean sensitivity 92.2%, mean specificity 86.0%) slightly dropped, but still SL performed better than any of the individual nodes (p<0.01 for swarm vs. each node, **Fig. 3b**). To determine whether sample size impacts on prediction results in this scenario, we reduced the number of samples at each training node (1-3) by 50%, but kept the ratio between cases and controls (**Extended Data Fig. S7c**). Still, SL outperformed the nodes, but all statistical readouts (mean accuracy 86.5%, mean sensitivity 87.8%, mean specificity 84.8%) at all nodes and SL showed lower performance, following general observations of AI with better performance when increasing training data^26^. We next altered the scenario by dividing up the three nodes into six smaller nodes (**Fig. 3c**, samples per node reduced by half in comparison to **Fig. 3a-b**), a scenario that can be envisioned in the domain of medicine in many settings, for example if several smaller medical centers with less cases would join efforts (**Fig. 3d**). Clearly, each individual node performed worse, but for SL the results did not deteriorate (mean accuracy 89.2%, mean sensitivity 90.7%, mean specificity 88.2% with significant difference to each of the nodes in all performance measures, see **Supplementary Table 4**), again illustrating the strength of the joined learning effort, while completely respecting each individual node’s data privacy.

**Figure 3.**
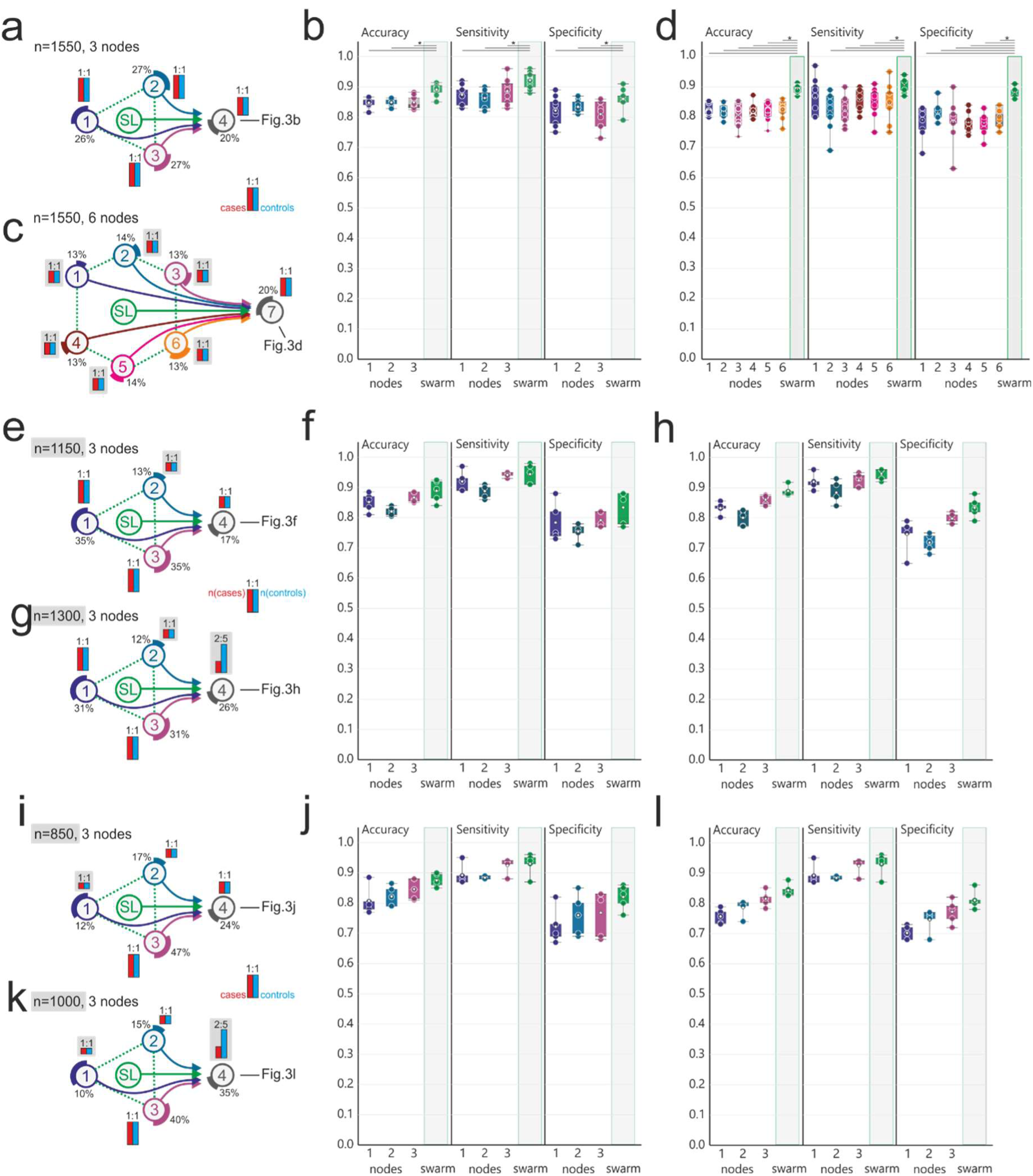
Swarm learning to identify patients with tuberculosis. **(a-l)** Principle of distribution of data to individual Swarm Edge Nodes (short ‘nodes’). Nodes 1-3 were used for training, node 4 for testing. Swarm Learning (SL) was achieved by integrating nodes 1-3 for training following procedures described in detail in Supplementary Information. All scenarios use dataset B and use acute TB as case and the remaining samples as controls. Left panels **(a,c,e,g,i,k)** illustrate the setting of the scenarios concerning distribution of cases (red bar) and controls (blue bar) to individual nodes, as well as total number of samples used for the scenario. Percentage at each node reflects the use of samples out of the complete dataset. **(a)** Scenario with even number of cases at each training node and the test node. **(b)** Evaluation of the scenario presented in (a) showing accuracy, sensitivity and specificity of five permutations for each training node and SL at node 4 (test node) as box-whisker plot (mean, 1st and 3rd quartile, whisker type Min/Max). **(c)** Scenario similar to (a) but with six training nodes. **(d)** Evaluation of scenario (c) as described in (b) but for all six training nodes. **(e)** Scenario where the training nodes have evenly distributed numbers of cases and controls at each training node, but node 2 has lower numbers of samples. **(f)** Evaluation of scenario (e) as described in (b). **(g)** Scenario similar to (e) but with reduced prevalence at the test node. **(h)** Evaluation of scenario (g) as described in (b). **(i)** Scenario with even distribution of cases and controls at each training node, but node 1 only has a very small training set. The test set is evenly distributed. **(j)** Evaluation of scenario (i) as described in (b). **(k)** Scenario similar to (i) but with uneven distribution in the test node. **(l)** Evaluation of scenario (k) as described in (b). Statistical differences between results derived by SL and individual nodes including all permutations performed were calculated with Wilcoxon signed rank test with continuity correction; asterisk and line: p<0.05.

Albeit aware of the fact that - in general - acute Tb is an endemic disease and does not tend to develop towards a pandemic such as the current COVID-19 pandemics, we utilized the Tb blood transcriptomics dataset to simulate potential outbreak and epidemic scenarios to determine benefits, but also potential limitations of SL and how to address them (**Fig. 3e-l**). The first scenario reflects a situation in which three independent regions (simulated by the nodes), would already have sufficient but different numbers of disease cases. Furthermore, cases and controls were kept even at the test node (**Fig. 3e-f**). Overall, compared to the scenario described in **Fig. 3c**, results for the swarm were almost comparable (mean accuracy 89.0%, mean sensitivity 94.4%, mean specificity 83.4%), while the results for the node with the lowest number of cases and controls (node 2) dropped noticeable (mean accuracy 82.2%, mean sensitivity 88.8%, mean specificity 75.4%, **Fig. 3f**). When reducing the prevalence at the test node by increasing the number of controls (**Fig. 3g-h**), this effect was even more pronounced, while the performance of the swarm was almost unaffected (mean balanced accuracy 89.0%).

We decreased the number of cases at a second training node (node 1) (**Fig. 3i-l**), which clearly reduced test performance for this particular node (**Fig. 3i-j**), while test performance of the swarm was only slightly inferior to the prior scenario (mean balanced accuracy 87.5%, no significant difference to the prior scenario). Only when reducing the prevalence at the test node (**Fig. 3k-l**), we saw a further drop in mean specificity for the swarm (81.0%), while sensitivity stayed similarly high (93.0%). Finally, we further reduced the prevalence at two training nodes (node 2: 1:10; node 3: 1:5) as well as the test node (**Extended Data Fig. 8a-b**). Lowering the prevalence during training resulted in very poor test performance at these two nodes (balanced accuracy node 2: 59.8%, balanced accuracy node 3: 74.8%), while specificity was high (node 2: 98.4%, node 3: 93.8%). SL showed highest accuracy (mean balanced accuracy 86.26%) and F-statistics (90.0%) but was outperformed for sensitivity by node 1 (swarm: 80.0%, node1: 87.8%), which showed poor performance concerning specificity (swarm: 92.4%, node1: 84.8%). Vice versa, node 2 outperformed the swarm for specificity (98.4%), but showed very poor sensitivity (21.2%) (**Extended Data Fig. 8b**). When lowering prevalence at the test node (**Extended Data Fig. 8c-d**), it became clear that all performance parameters including the F1 statistics were more resistant for the swarm compared to individual nodes. Taken together, using whole blood transcriptomes instead of PBMC and acute Tb as the disease instead of leukemia, we present a second use case illustrating that Swarm Learning integrating several individual nodes outperforms each node. Furthermore, we gained initial insights into the potential of SL to be utilized in a disease outbreak scenario.

### Identification of COVID-19 patients in an outbreak scenario

Based on the promising results obtained for tuberculosis, we collected blood from COVID-19 patients at two sites in Europe (Athens, Greece; n=39 samples, Nijmegen, n=93 samples) and generated whole blood transcriptomes by RNA-sequencing. We used the dataset described for Tb as the framework and included the COVID-19 samples (**Fig. 1l**) for assessing whether SL could be applied early on to detect patients with a newly identified disease. While COVID-19 patients are currently identified by PCR-based assays to detect viral RNA^53^, we use this case as a proof-of-principle study to illustrate how SL could be used even very early on during an outbreak based on the patients’ immune response captured by analysis of the circulating immune cells in the blood. Here, blood transcriptomes only present a potential feature space to illustrate the performance of SL. Furthermore, assessing the specific host response, in addition to disease prediction, might be beneficial in situations for which the pathogen is unknown, specific pathogen tests not yet possible, and blood transcriptomics can contribute to the understanding of the host’s immune response^54^. Lastly, while we do not have the power yet, blood transcriptome-based machine learning might be used to predict severe COVID-19 cases, which cannot be done by viral testing alone.

COVID-19 induces very strong changes in peripheral blood transcriptomes^54^. Following our experience with the leukemia and tuberculosis use cases, we first tested classifier performance for evenly distributed cases and controls at both training nodes and the test node (**Extended Data Fig. 9a,b, Supplementary Table 1)**. We reached very high statistical performance parameters, including high F1-statistics with SL showing highest mean values for accuracy (96.4%), sensitivity (97.8%), and F1 score (96.4%) (**Extended Data Fig. 9b**, summary statistics for all figures are listed in **Supplementary Table 4**). Reducing the prevalence at the test node (11:25 cases:controls) reduced all test parameters (**Extended Data Fig. 9c**), but only when we reduced the prevalence even further (1:44 ratio, **Extended Data Fig. 9d**), F1-statistics was clearly reduced, albeit SL again performing best. We next reduced the cases at all training nodes (**Extended Figure 10**), but even under these conditions, we observed still very high values for accuracy, sensitivity, specificity and F1 scores, both derived by training at individual nodes or by SL (**Extended Figure 10a-f**).

We then reduced the cases at all three training nodes to very low numbers, a scenario that might be envisioned very early during an outbreak scenario (**Fig. 4a**). Node 1 contained only 20 cases, node 2 10 cases and node 3 only 5 cases. At each node, controls outnumbered cases by 1:5, 1:10, or 1:20. At the test node, we varied the prevalence from 1:1 (**Fig. 4b**), 1:2 (**Fig. 4c**) to 1:10 (**Fig. 4d**). Based on our findings for Tb (**Extended Data Fig. 8**), we expected classifier performance to deteriorate under these conditions. We only observed decreased performance at nodes 2 and 3 in these scenarios with SL outperforming these nodes with p<0.05 for all performance measures, e.g. at a test node prevalence of 1:10 (accuracy (99.3%), sensitivity (95.1%), specificity (99.7%) and F1-statistics (99.7%) (**Fig. 4d**). Finally, we simulated a scenario with four instead of three training nodes with very few cases per node (**Extended Data Fig. 11a-d**), in an otherwise similar scenario as described for Fig. 4. Even for a simulated prevalence of 1:10 cases versus controls at the test node, we determined high test performance parameters for SL, with swam performance being significantly higher than node performances (SL accuracy (99.1%), sensitivity (92.0%), specificity (99.9%), F1 statistics (99.7%) (**Extended Data Fig. 11**) with the lowest variance in performance, while results at individual notes were very variable and deteriorated with low case numbers at the training node. Collectively, we provide first evidence that blood transcriptomes taken from patients with COVID-19 harbor very strong biological changes and these translate into a very powerful feature space for applying machine learning to the detection of patients with this new infectious disease, particularly when applying SL.

**Figure 4.**
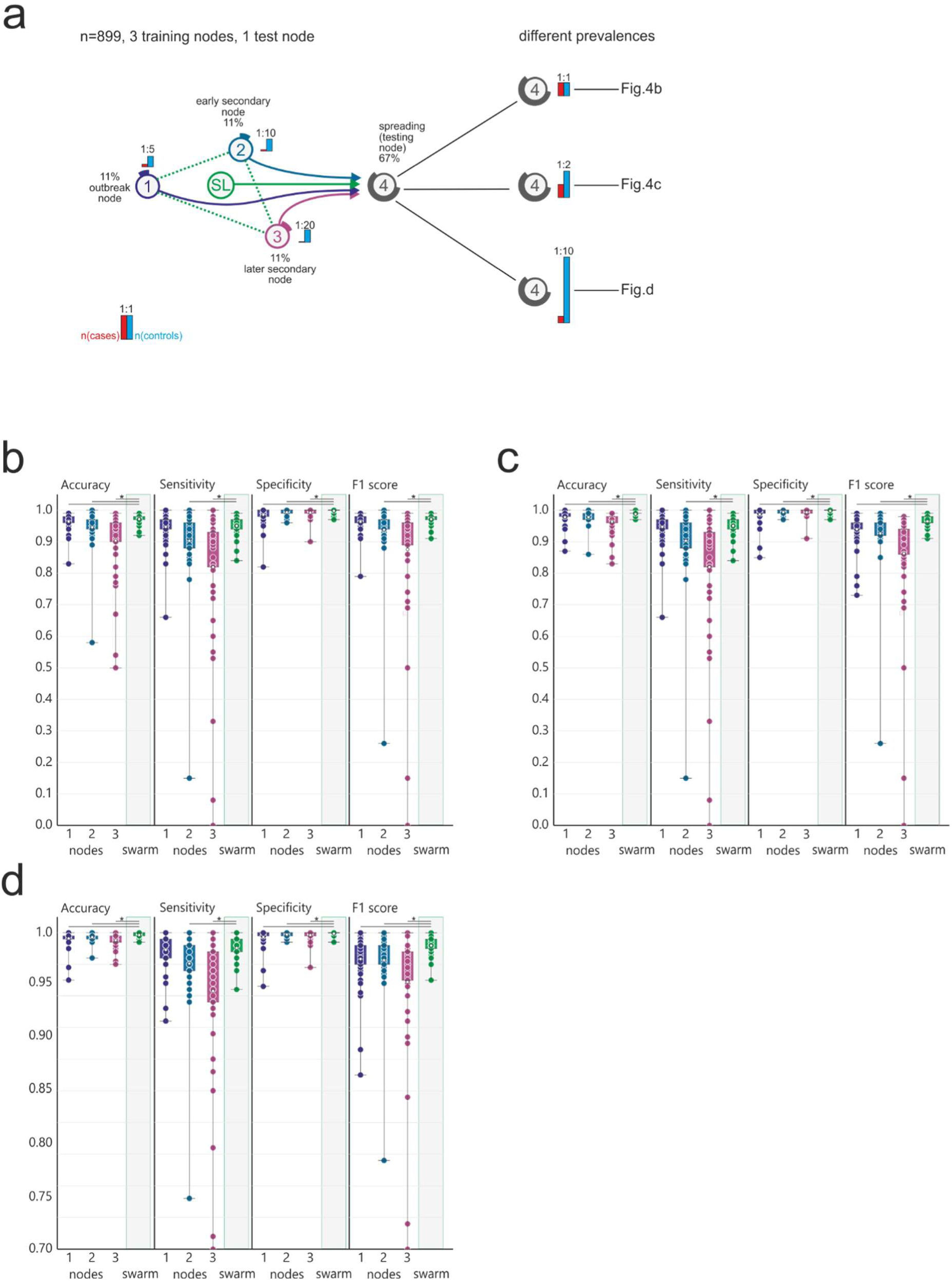
Identification of COVID-19 patients in an outbreak scenario. **(a)** Description of an outbreak scenario for COVID-19 using Dataset C. Nodes 1-3 were used for training, node 4 for testing. Swarm Learning (SL) was achieved by integrating nodes 1-3 for training following procedures described in detail in Supplementary Information. COVID-19 samples were used as cases. In this scenario, node 1 would be the outbreak node with the highest prevalence. Training node 2 has fewer cases and is an early secondary node, and node 3 acts as a later secondary node. The spreading is tested on the testing node with three different prevalences (**b,c,d**) and shown as box-whisker plot (mean, 1st and 3rd quartile, whisker type Min/Max). **(b)** Evaluation of (a) with even prevalence showing accuracy, sensitivity, specificity and F1-score of fifty permutations for each training node and the SL (node 4). **(c)** Evaluation (as described in (b)) of scenario (a) using a 1:2 ratio for cases and controls in the test set. **(d)** Evaluation (as described in (b)) of scenario (a) using a 1:10 ratio in the test set to simulate detection in regions with new infections. Statistical differences between results derived by SL and individual nodes including all permutations performed were calculated with Wilcoxon signed rank test with continuity correction; asterisk and line: p<0.05.

## Discussion

The introduction of precision medicine based on high-resolution molecular and imaging data will heavily rely on trustworthy machine learning algorithms in compute environments that are characterized by high accuracy and efficiency, that are privacy- and ethics-preserving, secure, and that are fault-tolerant by design^33–36^. At the same time, privacy legislation is becoming increasingly strict, as risks of cloud-based and central data-acquisition are recognized. Here, we introduce Swarm Learning, which combines blockchain technology and machine learning environments organized in a swarm network architecture with independent swarm edge nodes that harbor local data, compute infrastructure and execute the shared learning models that make central data acquisition obsolete. During iterations of SL, one of the nodes is chosen to lead the iteration, which does not require a central parameter server anymore thereby restricting centralization of learned knowledge and at the same time increasing resiliency and fault tolerance. In fact, these are the most important improvements over current federated computing models. Furthermore, private permissioned blockchain technology, harboring all rules of interaction between the nodes, is the Swarm Learning’s inherent privacy- and ethics-preserving strategy. This is of particular interest to medical data and could be adapted by other federated learning systems. To understand whether the concept of swarm learning would also be characterized by high efficiency and high accuracy, we built three medical use cases based on blood transcriptome data, which are high-dimensional data derived from blood, one of the major tissues used for diagnostic purposes in medicine. First, utilizing three previously compiled datasets (A1-3) of peripheral blood mononuclear cells derived from patients with acute myeloid leukemia, we provide strong evidence that SL-based classifier generation using a well-established neural network algorithm outperforms individual nodes, even in scenarios where individual contributing swarm nodes were performing rather poorly. Most striking, swarm learning was even improving performance parameters when training of individual nodes was based on technically different data, a situation that was previously shown to deteriorate classifier performance^47^. With these promising results, we generated a more challenging use case in infectious disease patients, detecting Tb based on full blood transcriptomes. Also in this case, SL outperformed individual nodes. Using Tb to simulate scenarios that could be envisioned for building blood transcriptome classifiers for patients during an outbreak situation, we further illustrate the power of SL over individual nodes. Considering the difficulty to quickly negotiate data sharing protocols or contracts during an epidemic or pandemic outbreak, we deduce from these findings that SL would be an ideal strategy for independent producer of medical data to quickly team up to increase the power to generate robust and reliable machine learning-based disease or outcome prediction classifier without the need to share data or relocate data to central cloud storages.

In addition, we tested whether we could build a disease prediction classifier for COVID-19 in an outbreak scenario. Building on our knowledge that blood transcriptomes of COVID-19 patients are significantly altered with hundreds of genes being changed in expression and with a rather specific signature compared to other infectious diseases^54^, we hypothesized that it should be possible to build such a classifier with a rather small number of samples. Here, we provide evidence that classifiers with high accuracy, sensitivity, specificity, and also high F1-statistics can be generated to identify patients with COVID-19 based on their blood transcriptomes. Moreover, we illustrate the power of SL that would allow to quickly increase the power of classifier generation even under very early outbreak scenarios with very few cases used at the training nodes, which could be e.g. collaborating hospitals in an outbreak region. Since data do not have to be shared, additional hospitals could benefit from such a system by applying the classifiers to their new patients and once classified, one could even envision an onboarding of these hospitals for an adaptive classifier improvement schema. Albeit technically feasible, we are fully aware that such scenarios require further classifier testing and confirmation, but also an assessment of how this could be integrated in existing legal and ethical regulations at different regions in the world^5, 6^. Furthermore, we appreciate that other currently less expensive data might be suitable for generating classifiers to identify COVID-19 patients^10^. For example, if highly standardized clinical data would become available, SL could be used to interrogate the clinical feature space at many clinics worldwide without any need to exchange the data to develop high performance classifiers for detecting COVID-19 patients. Similarly, recently introduced AI-systems using imaging data^21, 22^ might be more easily scaled if many hospitals with such data could be connected via SL. Irrespective of these additional opportunities using other parameter spaces, we would like to suggest blood transcriptomics as a promising new alternative due to its very strong signal in COVID-19. A next step will be to determine whether blood transcriptomes taken at early time points could be used to predict severe disease courses, which might allow physicians to introduce novel treatments at an earlier time point. Furthermore, we propose to develop an international database of blood transcriptomes that could be utilized for the development of predictive classifiers in other infectious and non-infectious diseases as well. It could be envisioned that such an SL-based learning scheme could be deployed as a permanent monitoring or early warning system that runs by default, looking for unusual movements in molecular profiles. Collectively, SL together with transcriptomics but also other medical data is a very promising approach to democratize the use of AI among the many stakeholders in the domain of medicine while at the same time resulting in more data privacy, data protection and less data traffic.

With increasing efforts to enforce data privacy and security of medical data^8^ (hhs.gov, https://www.hhs.gov/hipaa/index.html, 2020; Intersoft Consulting, General Data Protection Regulation, https://gdpr-info.eu) and to reduce data traffic and duplication of large medical data, a decentralized data model will become the preferred choice of handling, storing, managing and analyzing medical data^26^. This will not be restricted to omics data as exemplified here, but will extend to other large medical data such as medical imaging data^55, 56^. Particularly in oncology, great successes applying machine learning have already been reported for tumor detection^47, 55, 57, 58^, subtyping^59, 60^, grading^61^, genomic characterization^62^, or outcome prediction^63^, yet progress is hindered by too small datasets at any given institution^26^ with current privacy regulations^8^ (hhs.gov, https://www.hhs.gov/hipaa/index.html, 2020; Intersoft Consulting, General Data Protection Regulation, https://gdpr-info.ee) making it less appealing to develop centralized AI systems. We introduce Swarm Learning as a decentralized learning system with access to data stored locally that can replace the current paradigm of data sharing and centralized storage while preserving data privacy in cross-institutional research in a wide spectrum of biomedical disciplines. Furthermore, SL can easily inherit developments to further preserve privacy such as functional encryption^64^, or encrypted transfer learning approaches^65^. In addition, the blockchain technology applied here provides robust measures against semi-honest or dishonest participants/adversaries who might attempt to undermine a Swarm Network. Another important aspect for wide employment of SL in the research community and in real-world applications is the ease of use of the Swarm API, which will make it easier for researchers and developers to include novel developments such as for example private machine learning in TensorFlow^66^.

There is no doubt that numerous medical and other data types as well as a vast variety of computational approaches can be used during a pandemic^14^. We do not want to imply that blood transcriptomics would be the preferred solution for the many questions that AI and machine learning could help to solve during such a crisis. Although, at the same time, we have recently shown that blood transcriptomics can be used to define molecular phenotypes of COVID-19, uncover the deviated immune response in severe COVID-19 patients, define unique patterns of the disease in comparison to other diseases and can be utilized to predict potential drugs to be repurposed for COVID-19 therapy (Aschenbrenner et al. unpublished results). Therefore, we explored blood transcriptomics as a unique and rich feature space and a good example to illustrate the advantages of SL in identifying COVID-19 patients. Once larger datasets become available, SL could be used to identify patients at risk to develop severe COVID-19 early after onset of symptoms.

Another important quest that has been proposed is global collaboration and data-sharing^13^. While we could not agree more about the need for global collaboration - an inherent characteristic of SL - we favor systems that do not require data sharing but rather support global collaboration with complete data privacy preservation. Particularly, if using medical data that can also be used to interrogate medical issues unrelated to COVID-19. Indeed, statements by lawmakers have been triggered, clearly indicating that privacy rules also fully apply during the pandemics (EU Digital Solidarity: a call for a pan-European approach against the pandemic, Wojciech Wiewiórowski, https://edps.europa.eu/sites/edp/files/publication/2020-04-06_eu_digital_solidarity_covid19_en.pdf, 2020). Particular in a crisis situation such as the current pandemic, AI systems need to comply with ethical principles and respect human rights^14^. We therefore argue that systems such as Swarm Learning that allow fair, transparent and still highly regulated shared data analytics while preserving data privacy regulations are to be favored, particularly during times of high urgency to develop supportive tools for medical decision making. We therefore also propose to explore SL for image-based diagnostics of COVID-19 from patterns in X-ray images or computed tomography (CT) scans^21, 22^, structured health records^67^, or wearables for disease tracking^14^. Swarm learning would also have the advantage that model and code sharing as well as dissemination of new applications is easily scalable, because onboarding of new swarm participants is structured by blockchain technology, while scaling of data sharing is not even necessary due the inherent local computing of the data^14^. Furthermore, swarm learning can reduce the burden of establishing global, comprehensive, open, and verified datasets.

Collectively, we introduce Swarm Learning defined by the combination of blockchain technology and decentralized machine learning in an entirely democratized approach eliminating a central player and therefore representing a uniquely fitting strategy for the inherently locally organized domain of medicine. We used blood transcriptomes in three scenarios as use cases since they combine blood as the most widely used surrogate tissue for diagnostic purposes with an omics technology producing high-dimensional data with many parameters. Since the deployment of Swarm Learning due to ease of use of Swarm Learning libraries is a rather simple task, we propose to expand the use of this technology and further develop such classifiers in a unifying fashion across centers worldwide without any need to share the data itself. Our use cases are supposed to serve as examples for other high-dimensional data in the domain of medicine, but certainly also many other areas of research and application against the pandemics and beyond.

## Acknowledgments

We thank Dr. Sach Mukherjee (Statistics and Machine Learning, German Center for Neurodegenerative Diseases) for feedback on the manuscript.

## Deutsche COVID-19 Omics Initiative (DeCOI)

Robert Bals, Alexander Bartholomäus, Anke Becker, Ezio Bonifacio, Peer Bork, Thomas Clavel, Maria Colome-Tatche, Andreas Diefenbach, Alexander Dilthey, Nicole Fischer, Konrad Förstner, Julien Gagneur, Alexander Goesmann, Torsten Hain, Michael Hummel, Stefan Janssen, René Kallies, Birte Kehr, Andreas Keller, Sarah Kim-Hellmuth, Christoph Klein, Oliver Kohlbacher, Jan Korbel, Ingo Kurth, Markus Landthaler, Yang Li, Kerstin Ludwig, Oliwia Makarewicz, Manja Marz, Alice McHardy, Christian Mertes, Markus Nöthen, Peter Nürnberg, Uwe Ohler, Stephan Ossowski, Jörg Overmann, Klaus Pfeffer, Alfred Pühler, Nikolaus Rajewsky, Markus Ralser, Olaf Rieß, Stephan Ripke, Ulisses Nunes da Rocha, Philip Rosenstiel, Antoine-Emmanuel Saliba, Leif Erik Sander, Birgit Sawitzki, Philipp Schiffer, Joachim L. Schultze, Alexander Sczyrba, Oliver Stegle, Jens Stoye, Fabian Theis, Janne Vehreschild, Jörg Vogel, Max von Kleist, Andreas Walker, Jörn Walter, Dagmar Wieczorek, John Ziebuhr

## Author contributions

Conceptualization, J.L.S; K.L.S, S.Ma.; Methodology, S.W.-H., C.S., R.S., M.D., B.M., C.M.S.; Software: M.L.; V.G, C.S., S.Ma., S.Mu.; Investigation, S.W.-H., K.S., M.K., A.D., M.N., H.T., T.U.; Biospecimen/ enzyme resources, M.K., P.P., M.G.N., S.K., E.G.-B., M.M.B.BM., C.K., M.G.; Writing – Original Draft, S.W.-H., H.S., K.L.S., M.B., J.L.S.; Writing – Review & Editing, S.W.-H., H.S., K.L.S., D.S., A.C.A., M.K., P.P., M.G.N., M.B., S.C., M.S.W., E.L.G., J.L.S.; Visualization, H.S., J.L.S; Supervision, H.S., K.L.S., J.L.S.; Project Administration, H.S., J.L.S.; Funding Acquisition, W.F., J.L.S., F.J.T., J.H., N.Y., A.K.S., M.G., and M.M.B.B.

---

**Extended Data Figure 1.**
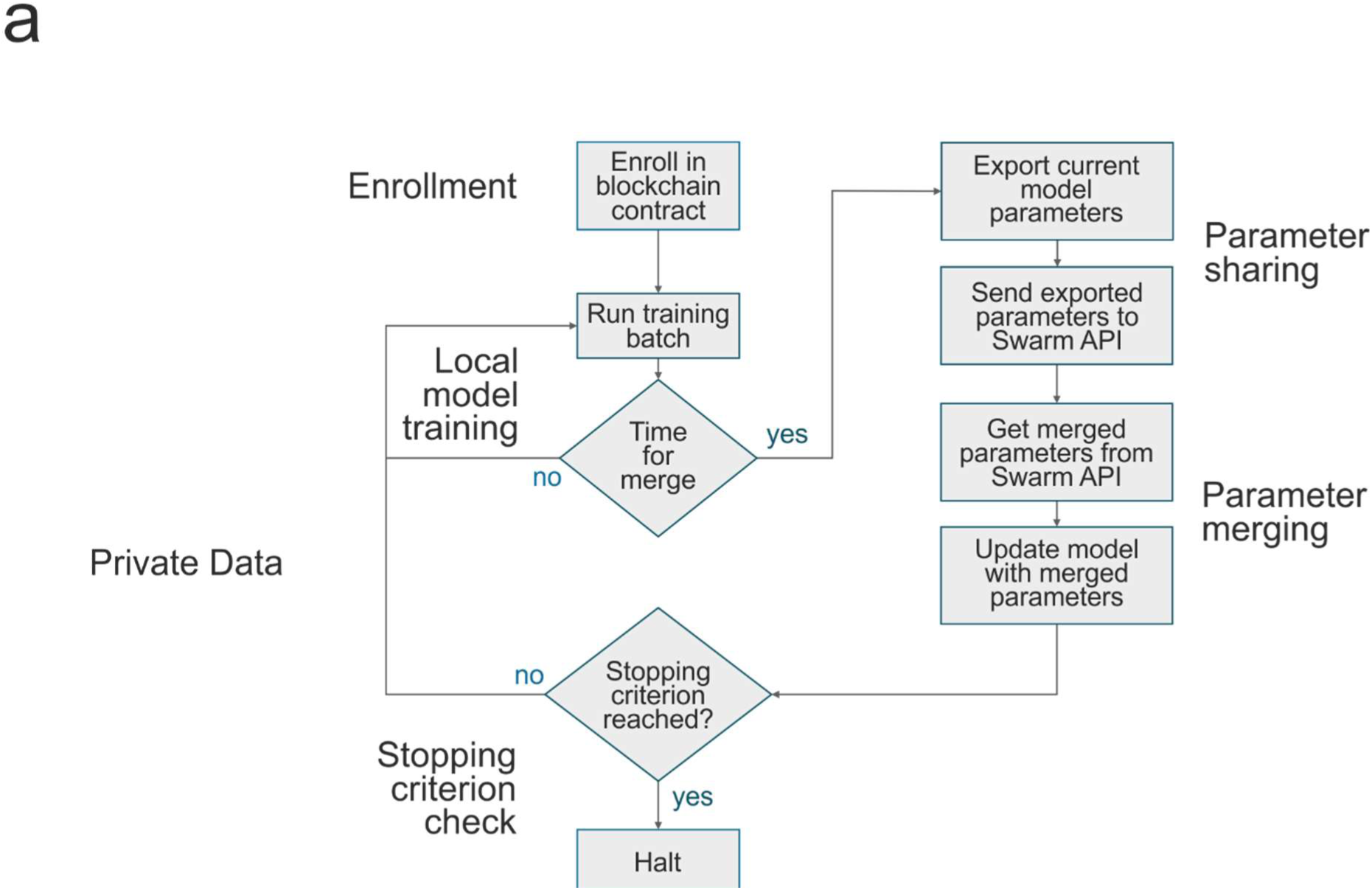
corresponding to Fig. 1. Schematics of the principles of the workflow of Swarm Learning once the nodes have been enrolled within the Swarm Network via private permissioned blockchain contract.

**Extended Data Figure 2.**
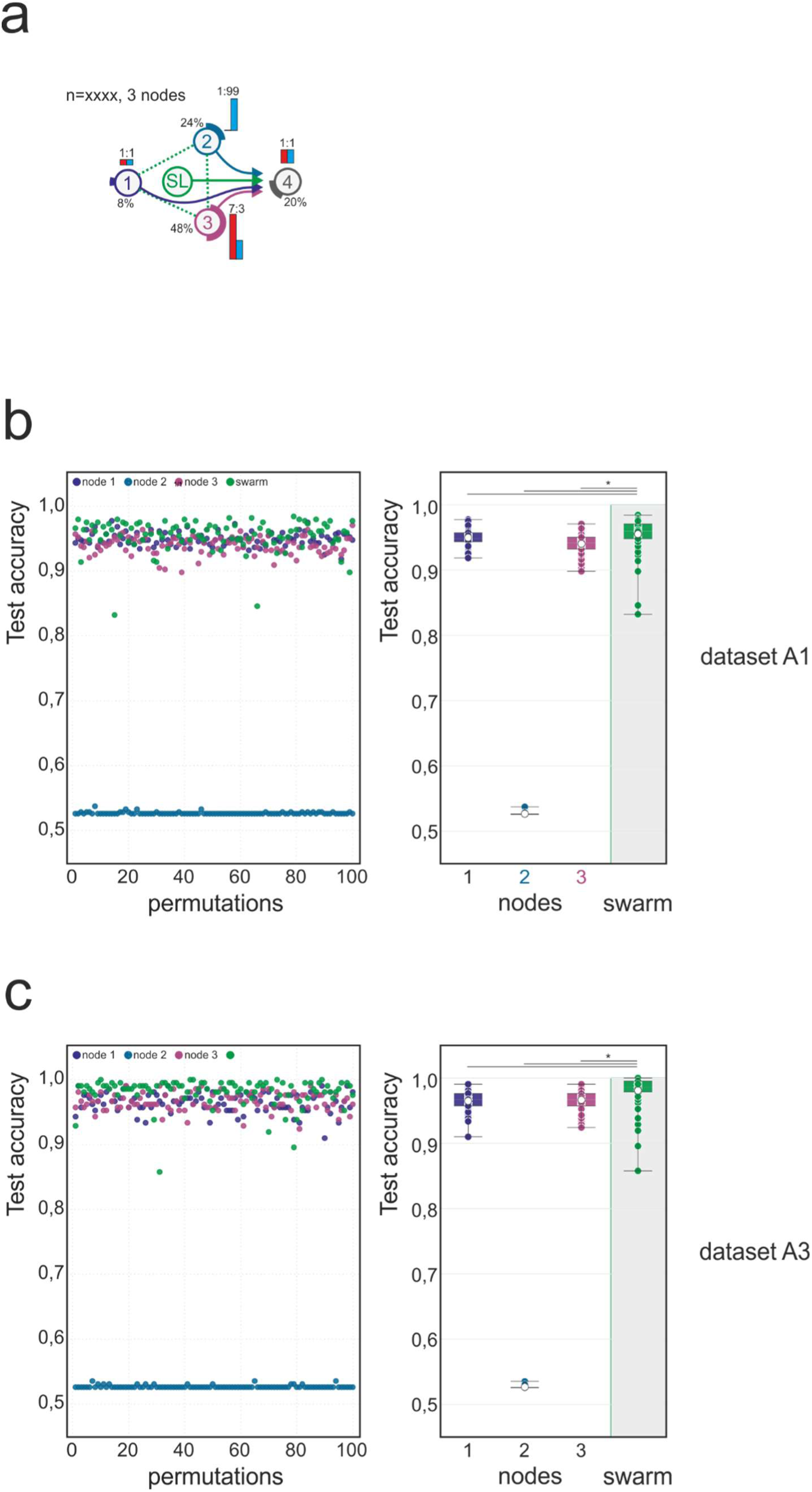
Scenario corresponding to Fig. 2b in dataset A1 and A3. Main settings are identical to what is described in Fig. 2 for Dataset A2. **(a)** Scenario with different prevalence of AML and different number of samples at each training node. The test set has an even distribution. **(b)** Evaluation of test accuracy for 100 permutations of dataset A1 per node and swarm. **(c)** Evaluation using dataset A3. Statistical differences between results derived by SL and individual nodes including all permutations performed were calculated with Wilcoxon signed rank test with continuity correction; asterisk and line: p<0.05.

**Extended Data Figure 3.**
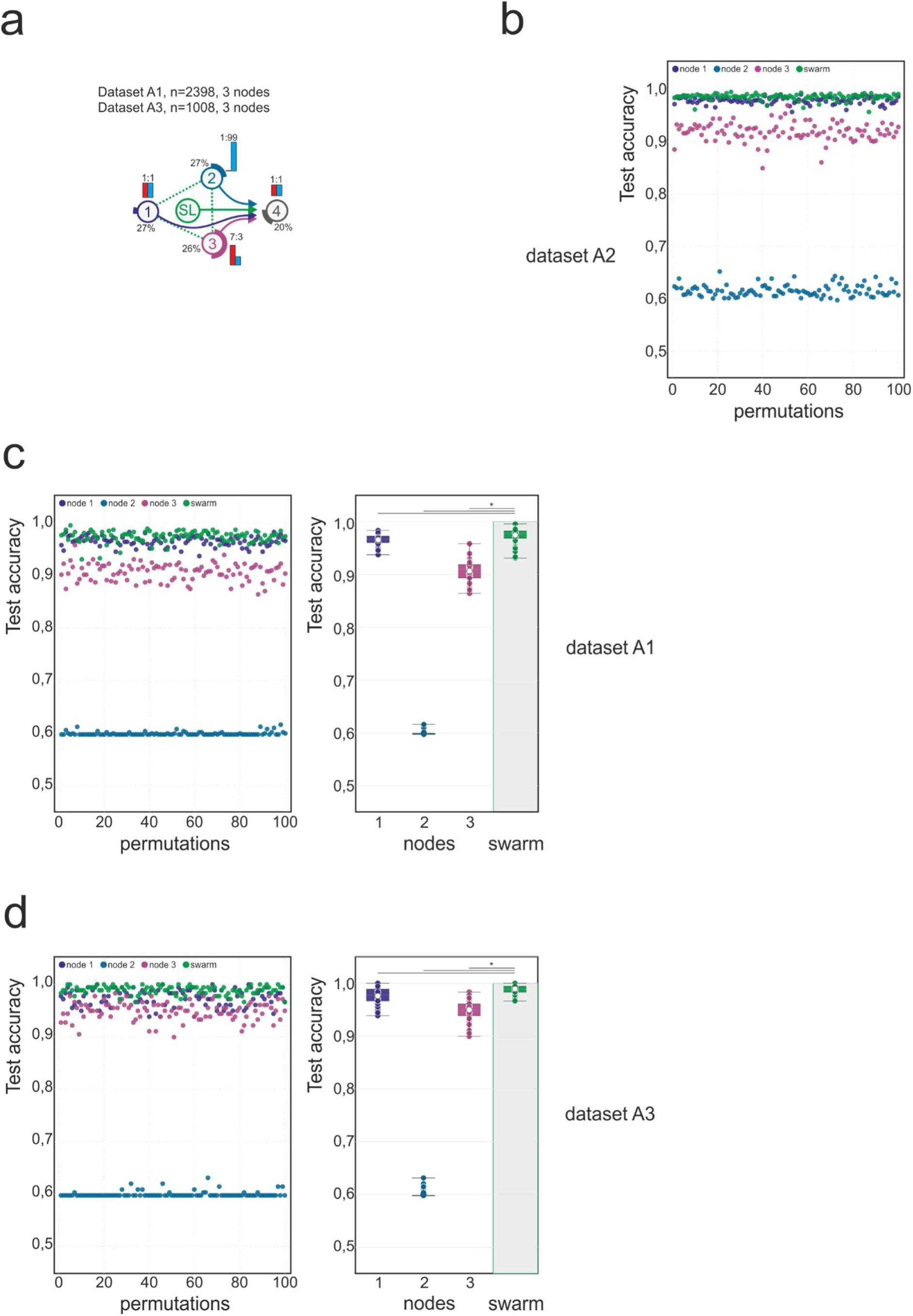
Scenario corresponding to Fig. 2c in dataset A1 and A3. Main settings are identical to what is described in Fig. 2 for dataset A2. **(a)** Scenario with similar training set sizes per node but decreasing prevalence. The test set ratio is 1:1. **(b)** Evaluation of the test accuracy over 100 permutation for dataset A2 (corresponding to Fig. 2c). **(c)** Evaluation of the test accuracy over 100 permutation for dataset A1. **(d)** Evaluation of the test accuracy over 100 permutation for dataset A3. Box-whisker plots (mean, 1st and 3rd quartile, whisker type Min/Max). Statistical differences between results derived by SL and individual nodes including all permutations performed were calculated with Wilcoxon signed rank test with continuity correction; asterisk and line: p<0.05.

**Extended Data Figure 4.**
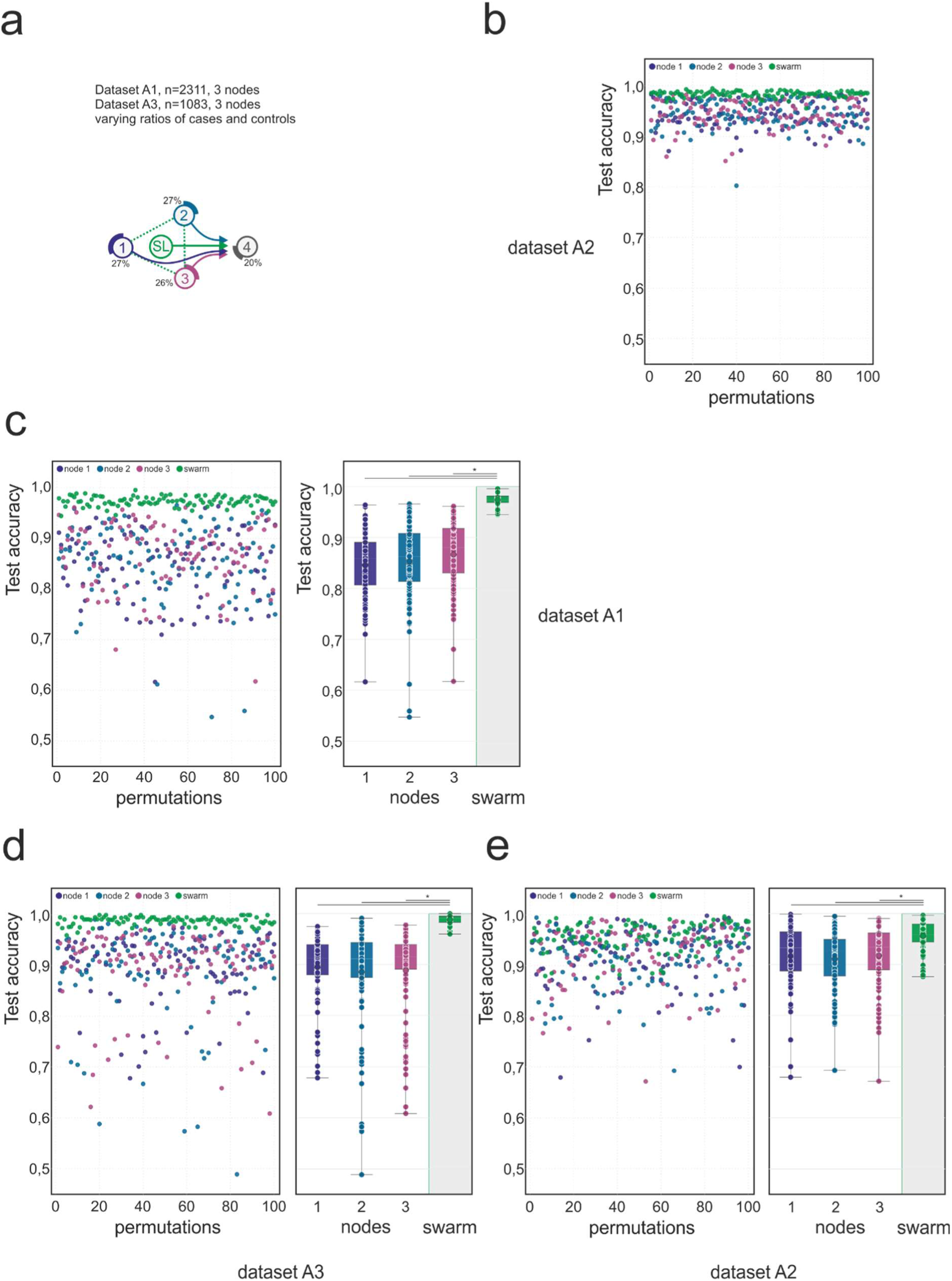
Scenario corresponding to Fig. 2d in dataset A1 and A3. Main settings are identical to what is described in Fig. 2 for dataset A2. (a) Scenario with similar sample sizes among three nodes, but with independent studies at each training node. Case and control ratios varied for each permutation. Testing samples are sampled from the studies also present in the training data. **(b)** Evaluation of the test accuracy over 100 permutation for dataset A2 (corresponding to Fig. 2d). **(c)** Evaluation of the test accuracy over 100 permutation for dataset A1. **(d)** Evaluation of the test accuracy over 100 permutation for dataset A3. **(e)** In this scenario, samples at the test node were derived from published studies completely independent from the studies used for training at the training nodes. Evaluation of the test accuracy over 100 permutation for dataset A2. Box-whisker plots (mean, 1st and 3rd quartile, whisker type Min/Max). Statistical differences between results derived by SL and individual nodes including all permutations performed were calculated with Wilcoxon signed rank test with continuity correction; asterisk and line: p<0.05.

**Extended Data Figure 5.**
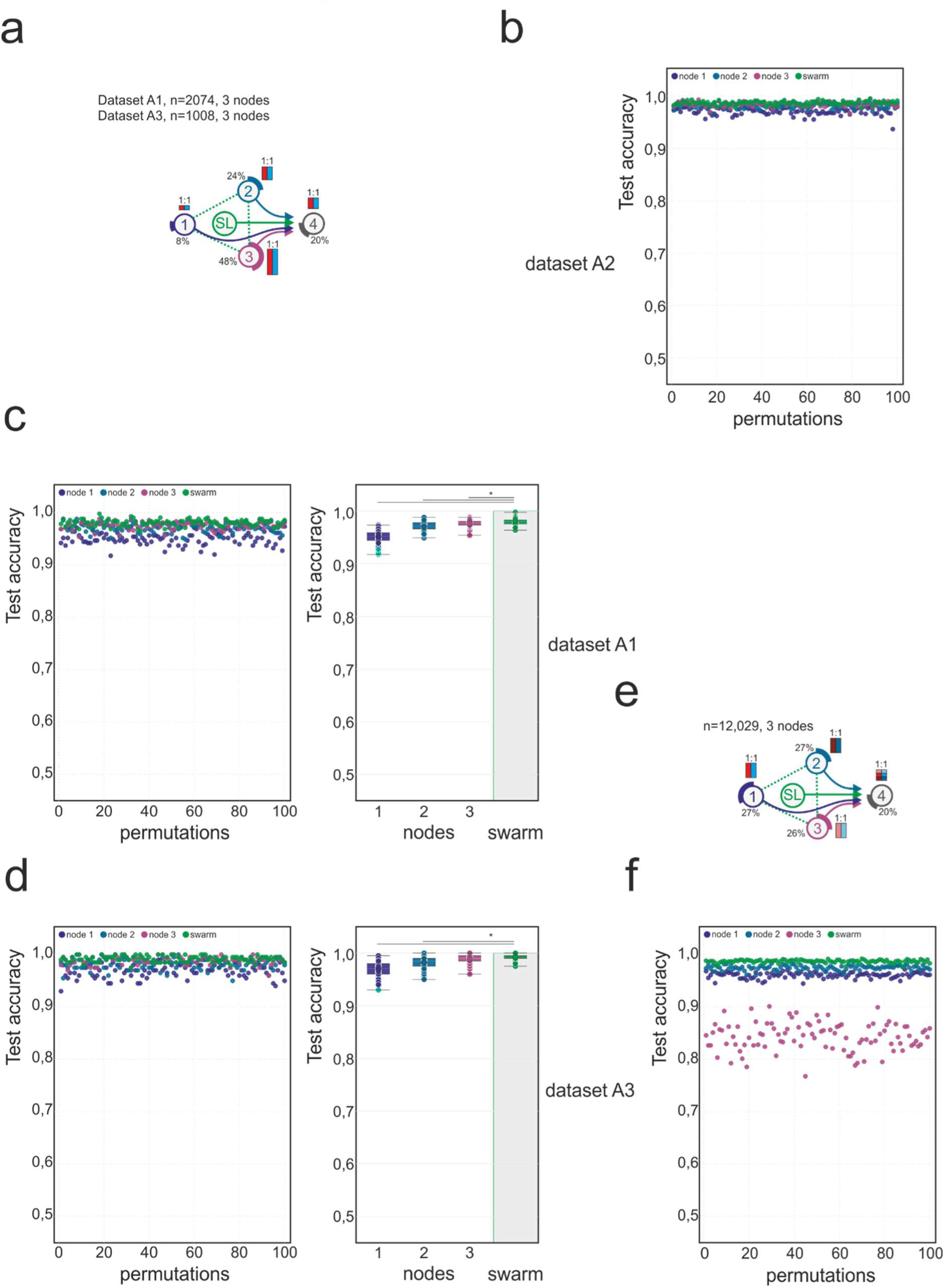
Scenario corresponding to Fig. 2e in dataset A1 and A3. Main settings are identical to what is described in Fig. 2 for dataset A2. **(a)** The case:control distribution is even, the training sets increase from node 1 to node 3. The test set is evenly split. **(b)** Test accuracy for evaluation of dataset A2 (corresponding to Fig. 2e). **(c)** Test accuracy for evaluation of dataset A1. **(d)** Test accuracy for evaluation of dataset A3. **(e)** Scenario where the data sets A1, A2, and A3 are assigned to a single training node each. Scenario similar to (a) but with equal training set sizes. **(f)** Evaluation results of 100 permutations (corresponding to Fig. 2f). Box-whisker plots (mean, 1st and 3rd quartile, whisker type Min/Max). Statistical differences between results derived by SL and individual nodes including all permutations performed were calculated with Wilcoxon signed rank test with continuity correction; asterisk and line: p<0.05.

**Extended Data Figure 6.**
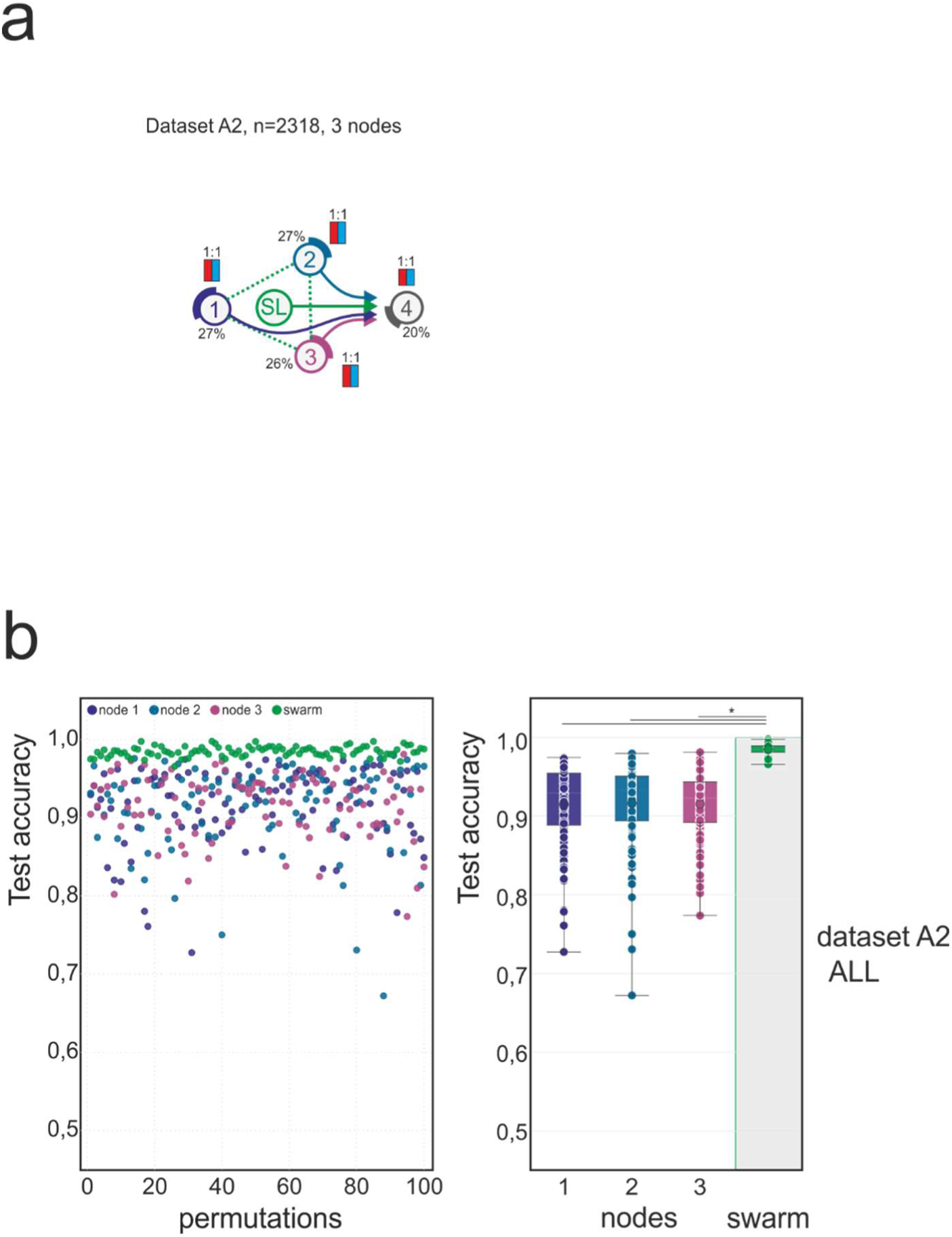
Scenario for ALL in dataset 2. Main settings are identical to what is described in Fig. 2 for dataset A2. Here cases are samples derived from patients with ALL, while all other samples are controls (including AML). **(a)** Scenario for the detection of ALL in dataset A2. The training sets are evenly distributed among the nodes. The test ratio is 1:1. **(b)** Evaluation of scenario (a) for test accuracy over 100 permutations. Box-whisker plot (mean, 1st and 3rd quartile, whisker type Min/Max). Statistical differences between results derived by SL and individual nodes including all permutations performed were calculated with Wilcoxon signed rank test with continuity correction; asterisk and line: p<0.05.

**Extended Data Figure 7.**
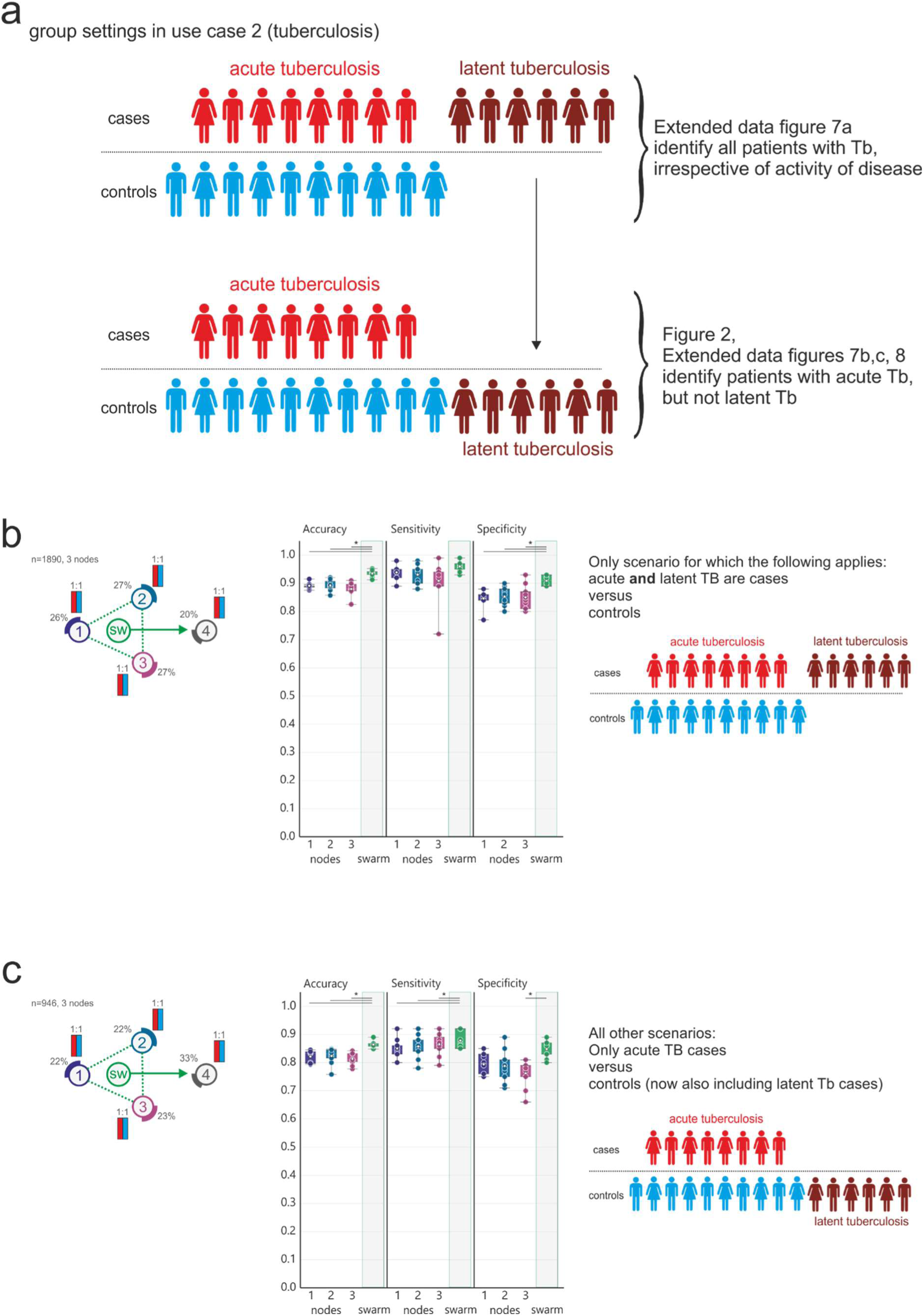
Scenario for detecting all Tb versus controls. **(a)** Description of the different group settings used based on the assignment of latent Tb to control or case. **(b)** Evaluation of a scenario where acute and latent Tb are cases. The data is evenly distributed among the training nodes. The scenario is evaluated as described in Figure 3 (b). **(c)** Scenario designed similar to (b) but latent Tb is part of control. Box-whisker plot (mean, 1st and 3rd quartile, whisker type Min/Max). Statistical differences between results derived by SL and individual nodes including all permutations performed were calculated with Wilcoxon signed rank test with continuity correction; asterisk and line: p<0.05.

**Extended Data Figure 8:**
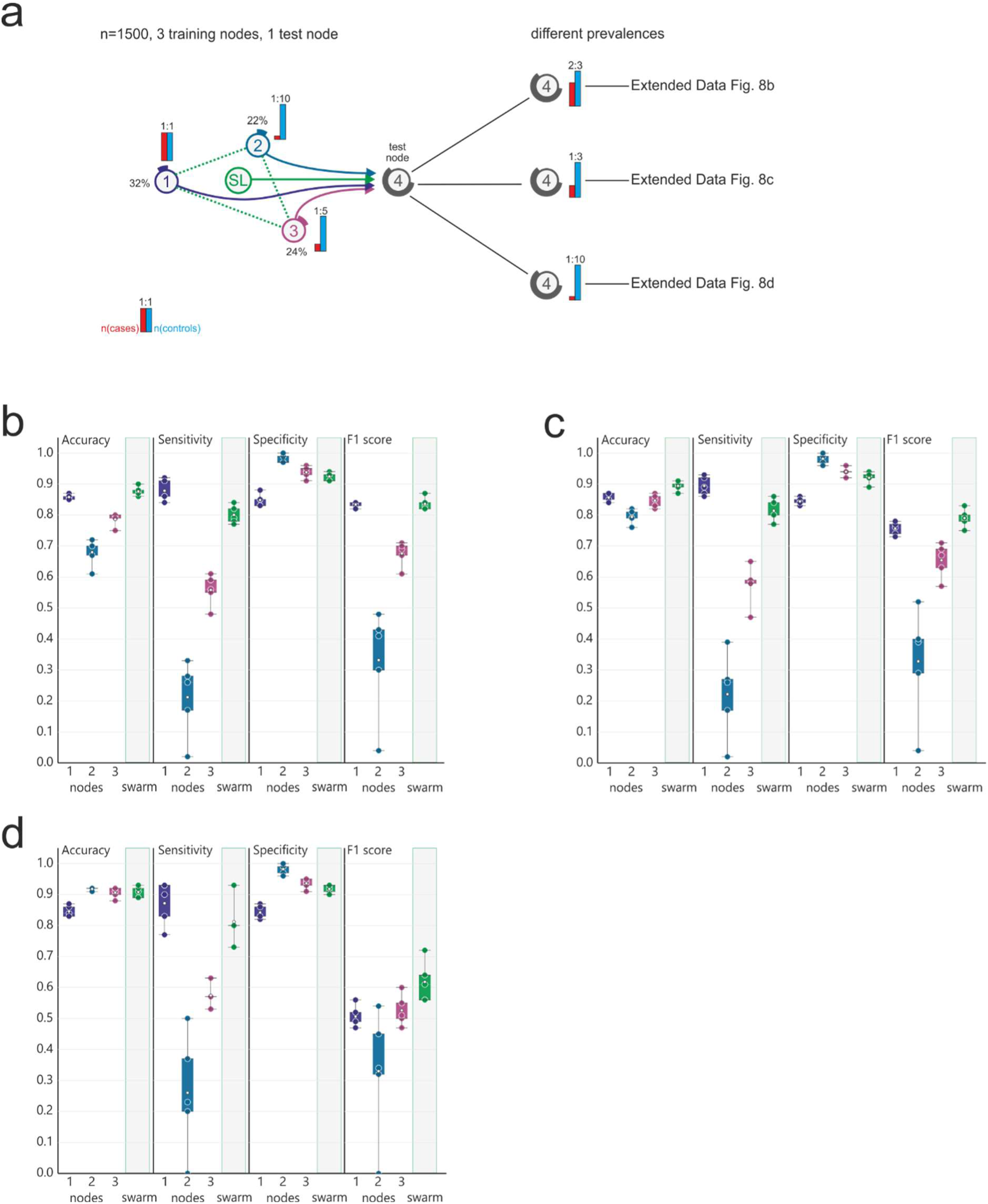
Scenario detecting acute Tb with low prevalence at training nodes. **(a)** Scenario with training nodes having different prevalence: node 2 has only a 1:10 ratio. Three prevalence scenarios are used in the test set. **(b)** Evaluation of scenario (a) showing accuracy, sensitivity, specificity and F1 score. **(c)** Similar scenario as in (a) but prevalence changed to 1:3 cases: controls in the training set. **(d)** Similar scenario as in (a) but prevalence changed to 1:10 cases: controls in the training set. Box-whisker plot (mean, 1st and 3rd quartile, whisker type Min/Max). Statistical differences between results derived by SL and individual nodes including all permutations performed were calculated with Wilcoxon signed rank test with continuity correction; asterisk and line: p<0.05.

**Extended Data Figure 9.**
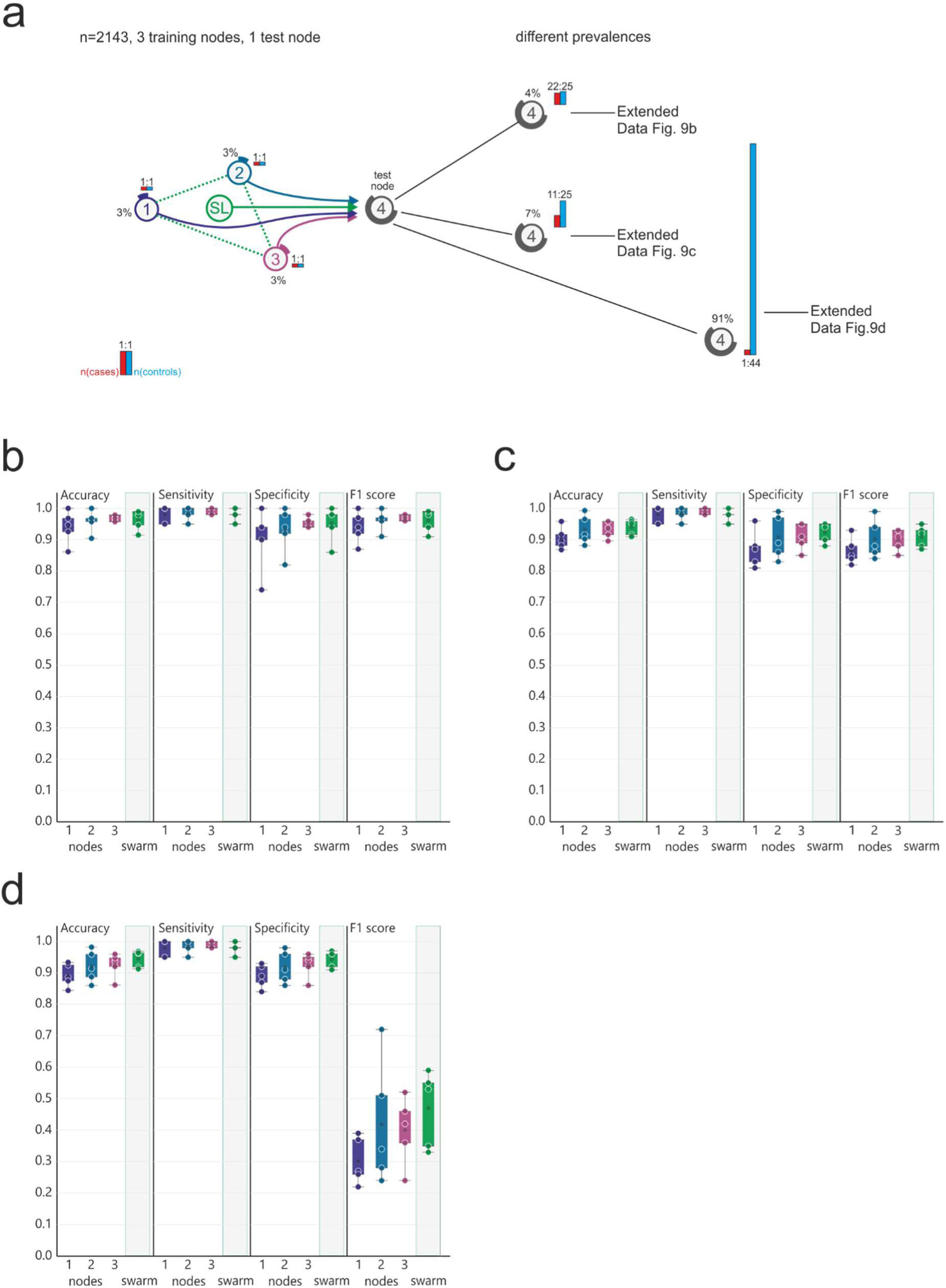
Baseline scenario for detecting COVID-19 patients. **(a)** Scenario with even training set distribution among nodes 1-3. Three different testing sets with different prevalence are simulated. **(b)** Evaluation of (a) for a 22:25 case: control ratio showing accuracy, sensitivity, specificity and F1 score. **(c)** Evaluation results of scenario (a) for a 11:25 ratio. **(d)** Evaluation results of scenario (a) for a 1:44 prevalence. Box-whisker plot (mean, 1st and 3rd quartile, whisker type Min/Max). Statistical differences between results derived by SL and individual nodes including all permutations performed were calculated with Wilcoxon signed rank test with continuity correction; asterisk and line: p<0.05.

**Extended Data Figure 10.**
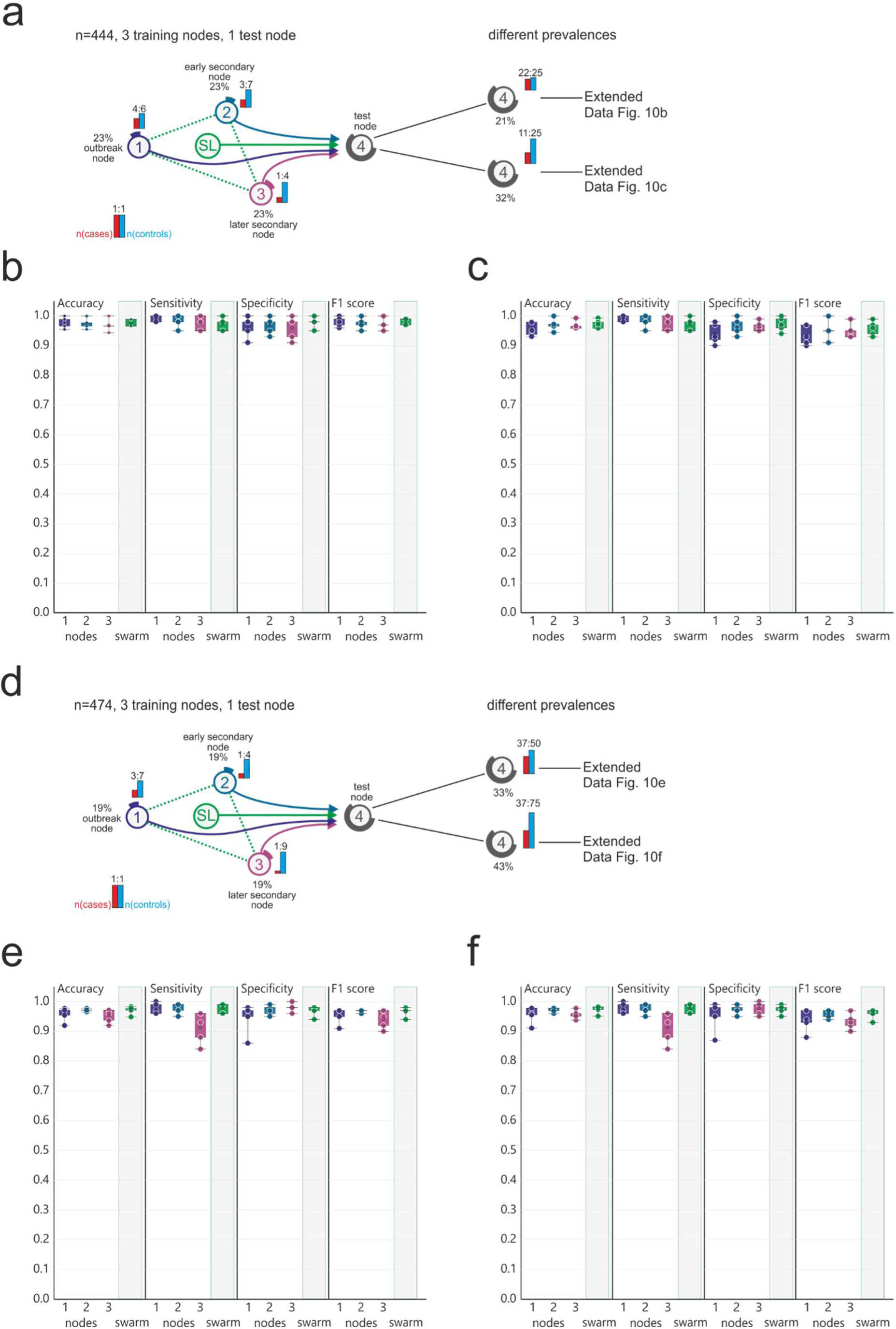
Scenario with reduced prevalence at training nodes for detecting COVID-19 patients. **(a)** This scenario has the same sample size at each training node, but the prevalence decreases from node 1 to node 3. There are two different test sets (b) and (c). **(b)** Evaluation of scenario (a) with 22:25 ratio at the test node. **(c)** Results for the evaluation of scenario (a) with reduced prevalence. **(d)** Scenario similar to (a) but the prevalence has a steeper decrease between node 1 and 3. **(e)** Evaluation of scenario (d) with a ratio of 37:50 at the test node. **(f)** Evaluation of (d) with a reduced prevalence compared to (e). Box-whisker plot (mean, 1st and 3rd quartile, whisker type Min/Max). Statistical differences between results derived by SL and individual nodes including all permutations performed were calculated with Wilcoxon signed rank test with continuity correction; asterisk and line: p<0.05.

**Extended Data Figure 11.**
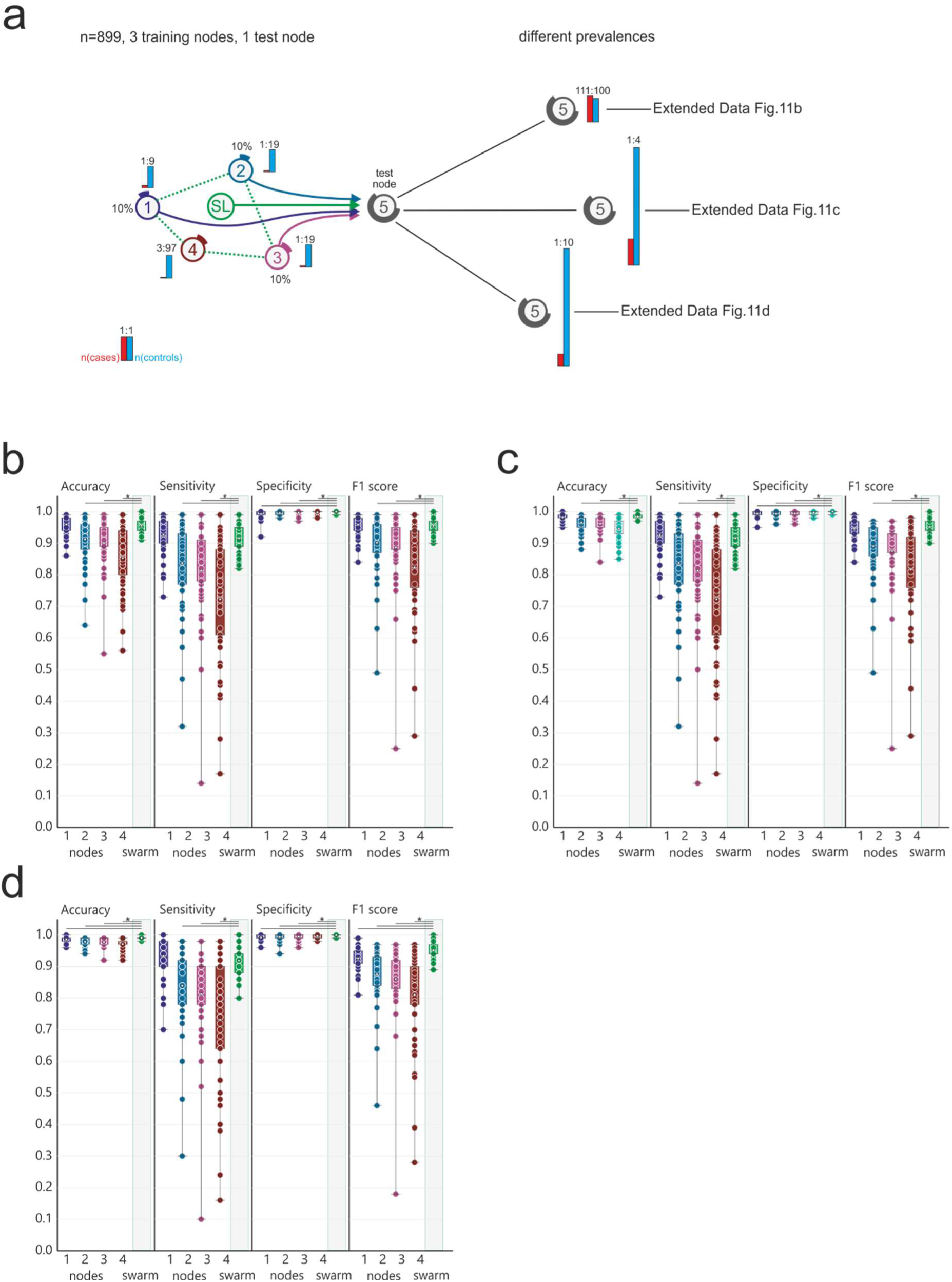
Scenario with reduced prevalence in training and test set at a 4-node setting. **(a)** This scenario has even training set sizes among the nodes with the prevalence ranging from 10% at node 1 to 3% at nodes 3 and 4. There are three different test sets (b), (c) and (d) with decreasing prevalence and increasing total sample size. **(b)** Evaluation of scenario (a) with 111:100 ratio. **(c)** Evaluation of scenario (a) with 1:4 ratio and increased sample number of the test set. **(d)** Results of scenario (a) with 1:10 prevalence and increased sample number of the test set. Box-whisker plot (mean, 1st and 3rd quartile, whisker type Min/Max). Statistical differences between results derived by SL and individual nodes including all permutations performed were calculated with Wilcoxon signed rank test with continuity correction; asterisk and line: p<0.05.

## Supplementary Information

### (Material and Methods)

#### Datasets

##### Peripheral blood mononuclear cell (PBMC) derived transcriptome dataset (Dataset A)

We used a previously described dataset containing over 12,000 transcriptomes derived from peripheral blood mononuclear cells (PBMC), deposited at the National Center for Biotechnology Information Gene Expression Omnibus^68^ (GEO) under SuperSeries GSE122517 or via the individual SubSeries GSE122505 (dataset 1), GSE122511 (dataset 2) and GSE122515 (dataset 3). Briefly, this dataset was generated by inspection of all publicly available datasets at GEO on September 20th, 2017. Inclusion criteria were cell type (PMBCs) and species (*Homo sapiens*). Existing GEO SuperSeries were excluded to avoid duplicated samples. According to data generation method, three datasets were established; dataset 1, generated with Affymetrix HG-U133 A microarrays (n=2,500), dataset 2 with Affymetrix HG-U133 2.0 microarrays (n=8,348), and dataset 3 with high-throughput RNA sequencing (RNA-seq)(n=1,181). Data were curated as previously described^47^. All sample information is listed in **Supplementary Table 2.**

##### Whole blood derived transcriptomes for the prediction of tuberculosis (Dataset B)

To establish a dataset based on whole blood transcriptomes we generated new data from healthy controls (Rhineland Study) and combined these with previously generated data that had been deposited in Gene Expression Omnibus (GEO). We screened for transcriptome datasets derived from human whole blood samples, which were collected using the PAXgene Blood RNA System. In total, nine independent datasets were selected to be included in the present study (GSE101705 (n=44); GSE107104 (n=33), GSE112087 (n=120), GSE128078 (n=99), GSE66573 (n=14), GSE79362 (n=355), GSE84076 (n=36); GSE89403 (n=914)). The newly generated 384 whole blood samples were sampled in context of the Rhineland Study led by the German Center for Neurodegenerative Diseases (DZNE), which is an extensive longitudinal study monitoring healthy individuals over 2 decades. Approval to undertake the Rhineland Study was obtained from the ethics committee of the University of Bonn, Medical Faculty. The study is carried out in accordance with the recommendations of the International Conference on Harmonization (ICH) Good Clinical Practice (GCP) standards (ICH-GCP). Written informed consent was obtained from all participants in accordance with the Declaration of Helsinki. Overnight fasting blood was collected from all participants, including a PAXgene® tube for RNA extraction and RNA-seq analysis. In total, Dataset B contained 1999 samples from patients with active tuberculosis (n=775), latent tuberculosis (n=277), fatigue (n=55), autoimmune diseases (n=68), HIV (n=16) and controls (n=808). Sample information is listed in **Supplementary Table 2**.

##### Whole blood derived transcriptome dataset for the prediction of COVID-19 (Dataset C)

To develop classifiers based on whole blood transcriptomes able to predict COVID-19 patients we collected an additional 134 PAXgene® tubes for RNA extraction and RNA-seq analysis from COVID-19 patients, of which 93 whole blood samples at the Intensive Care Unit of the Radboud University Medical Centre in Nijmegen, the Netherlands, and 41 samples were either collected at the Sotiria Athens General Hospital or the ATTIKON University General Hospital in Athens, Greece. For all COVID-19 patients, the study was carried out in accordance with the applicable rules concerning the review of research ethics committees and informed consent. All patients or legal representatives were informed about the study details and could decline to participate. COVID-19 was diagnosed by a positive SARS-CoV-2 RT-PCR test in nasopharyngeal or throat swabs and/or by typical chest CT-scan finding. Blood for RNA-seq analysis was sampled on day 0 to 11 after admission. In the cohort in Athens, blood samples from ten healthy donors who were tested negative on SARS-CoV-2 were included as controls. The newly generated samples from the COVID-19 patients and the controls from Athens were combined with dataset B (see above) to establish Dataset C. As a result, in addition to the 1999 samples derived from Dataset B, Dataset C included further 10 healthy controls and 134 dutch COVID-19 samples, which makes a total of 2,143 samples. Sample information is listed in **Supplementary Tables 2 and 6**.

#### Pre-processing

##### PBMC transcriptome dataset (Dataset A)

We used a previously published dataset compiled for predicting AML in blood transcriptomes derived from peripheral blood mononuclear cells (PBMC)^47^. Briefly, all raw data files were downloaded from GEO and the RNA-seq data was preprocessed using the kallisto aligner against the human reference genome gencode v27 (GRCh38.p10). For normalization, we considered all platforms independently, meaning that normalization was performed separately for the samples in Dataset A1, A2 and A3, respectively. Microarray data (Datasets A1 and A2) was normalized using the robust multichip average (RMA) expression measures^69^, as implemented in the R package affy^70^. RNA-seq data (Dataset A3) was normalized with the R package DESeq2 using standard parameters^71^. In order to keep the datasets comparable, data was filtered for genes annotated in all three datasets, which resulted in 12,708 genes. No filtering of low-expressed genes was performed. All scripts used in this study for pre-processing are provided as a docker container on Docker Hub (docker hub, https://hub.docker.com/r/schultzelab/aml_classifier).

##### Whole blood derived transcriptome datasets (Datasets B and C)

Since alignment of whole blood transcriptome data can be performed in numerous different ways, we re-aligned all downloaded and collected datasets which were 4.7 Terabyte in size and comprised a total of 7.8 Terabases, to the human reference genome gencode v33 (GRCh38.p13) and quantified transcript counts using STAR, an ultrafast universal RNA-seq aligner (version 2.7.3a) ^72^. For all samples in Datasets B and C, raw counts were imported using DESeqDataSetFromMatrix function and size factors for normalization were calculated using the DESeq function using standard parameters^71^. This was done separately for Dataset B and Dataset C. Since some of the samples were prepared with poly-A selection to enrich for protein-coding mRNAs, we filtered the complete dataset for protein-coding genes in order to ensure greater comparability across library preparation protocols. Furthermore, we excluded all ribosomal protein-coding genes, as well as mitochondrial genes and genes coding for hemoglobins, which resulted in 18,135 transcripts as the feature space in Dataset B and 19,358 transcripts in Dataset C. Furthermore, transcripts with an overall expression < 10 were excluded from further analysis. Other than that, no filtering of transcripts was performed. Prior to use in machine learning we performed a rank transformation to normality on both datasets B and C^73^. Briefly, transcript expression values were transformed from RNAseq counts to their respective ranks. This was done transcript-wise, meaning all transcript expression values per sample were given a rank based on ordering them from lowest to highest value. The rankings were then turned into quantiles and transformed via the inverse cumulative distribution function of the Normal distribution. This leads to all transcripts following the exact same distribution (that is, a standard Normal with a mean of 0 and a standard deviation of 1) across all samples

#### Methods details

##### Scenarios for prediction of AML

We previously demonstrated that ML on PBMC transcriptomes can be utilized to predict AML^47^. Based on this experience, we generated sample sets within three independent transcriptome datasets (dataset A1-A3, see above) to assess different scenarios in a three-node setting for training with a fourth node only used for testing. As indicated in Fig. 2, six scenarios with varying numbers of samples per node and varying ratios between cases and controls at each node where defined. For predicting AML, all samples derived from AML patients were classified as cases, while all other samples were labeled controls. When predicting ALL, all samples derived from ALL patients were classified as cases and all others as controls. For each scenario (Fig. 2) and each dataset we permuted the sample distribution 100 times, resulting in a total of 5,594 individual predictions. The different scenarios were chosen to address the influence of sample numbers per node, the case control ratio, study design-related batch effects, and transcriptome technologies used on classifier performance at the nodes, but more importantly on swarm learning performance. Sample distributions for all permutations within all scenarios are listed in **Supplementary Table 1**.

##### Scenarios for detecting patients with acute TB

In line with the experience we gained from the prediction of AML, we used dataset B to generate scenarios for the prediction of tuberculosis in various settings, again using different scenarios in a three-node setting for training with a fourth node only used for testing. In one scenario, all patients with tuberculosis (Tb) including patients with latent and acute Tb were treated as cases, while all others were defined as controls (**Extended Data Fig. 6b**). In all other scenarios, cases were restricted to acute Tb patients’ samples, while patients with latent Tb were defined as controls together with all other non-Tb samples. Here, the question to be answered is, whether the classifiers can identify patients with acute Tb and can distinguish them from latent Tb and other conditions.

In one scenario (**Fig. 3c-d**), we added three additional training nodes to test dependency of classifier performance by the number of nodes. As indicated in Fig. 3, three scenarios with varying numbers of samples per node and varying ratios between cases and controls at each node where defined. For scenarios described within **Fig. 3e,g** and **Fig. 3i,k**, we tested two prevalence scenarios in the test set. For each scenario (**Fig. 3**) we permuted the sample distribution 5-10 times, resulting in a total of 325 individual predictions. To mimic an outbreak scenario, we reduced cases also at the training nodes to determine the effects on Swarm Learning performance. Sample distributions for all permutations within all scenarios are listed in **Supplementary Table 1**.

##### Simulation of an outbreak scenario to detect COVID-19 patients

Based on the promising results obtained with tuberculosis, we next intended to simulate classifier building and testing for the prediction of COVID-19 in a SL setting. We used dataset B and added 144 additional samples, of which 139 samples were derived from COVID-19 patients (see above). We applied a three-node setting for training with a fourth node only used for testing.

In one scenario (**Extended Data Fig. 8**), we kept cases (n=30) and controls (n=30) evenly distributed among the three training nodes and tested three different prevalence scenarios at the test node (22:25; 11:25; 1:44). In a second scenario (**Extended Data Fig. 9a-c**) we changed the ratio of cases and controls at each node (node 1: 40:60, node 2: 30:70, node 3: 20:80) and tested two prevalence scenarios at the test node (22:25; 11:25). In a third scenario (**Extended Data Fig. 9a-c**) we further reduced the number of cases at the training nodes further (node 1: 30:70, node 2: 20:80, node 3: 10:90) and tested two prevalence scenarios at the test node (37:50; 37:75).

Lastly, we tested an outbreak scenario (**Fig. 4**) with very few cases at the outbreak node 1 (20:80), an early secondary node (10:90) and a later secondary node (5:95) and three prevalence scenarios at the test node (1:1, 1:2, 1:10), resulting in a total of 220 individual predictions Sample distributions for all permutations within all scenarios are listed in **Supplementary Table 1**.

##### Application layer

The application layer (see also **Fig. 1g**) consists of disease models for which definitions are given, which samples are cases and which samples are controls. For example, if the classifier is supposed to detect all patients with tuberculosis (Tb), the model includes patients with latent and acute tuberculosis as cases and all other samples as controls. However, if only patients with acute tuberculosis are intended to be detected as cases, the model is changed in that cases are now only patient samples derived from patients with acute Tb, while samples from patients with latent Tb are now treated as controls, similar to all other non-Tb samples. The cases and controls used for each scenario are given in the result section in more detail. For each mode, classifiers are generated by applying neural networks (for description see below)

#### Computation and analysis

##### Neural network algorithm

We leveraged a deep neural network with a sequential architecture as implemented in the keras library (Keras, https://keras.io/, 2015). Briefly, the neural network consists of one input layer, eight hidden layers and one output layer. The input layer is densely connected and consists of 256 nodes, a rectified linear unit activation function and a dropout rate of 40%. From the first to the eighth hidden layer, nodes are reduced from 1024 to 64 nodes, and all layers contain a rectified linear unit activation function, a kernel regularization with an L2 regularization factor of 0.005 and a dropout rate of 30%. The output layer is densely connected and consists of 1 node and a sigmoid activation function. The model is configured for training with Adam optimization and to compute the binary cross-entropy loss between true labels and predicted labels.

The model has been translated from R to Python in order to make it compatible with the swarm learning library. This model is used for training both the individual nodes as well as swarm learning. The model is trained over 100 epochs, with varying batch sizes. The batch size of 8, 16, 32, 64 and 128 are used depending on the number of training samples.

##### Preparation and adaptation of neural network code to be used in a swarm learning environment

A swarm callback is introduced to integrate the model with the Swarm Learning library. Minimum number of nodes for synchronization, synchronization interval, validation dataset and batch size are passed as parameters to swarm callback. The swarm call back API is

~~~
swCallback = SwarmCallback(sync_interval = <number of training batches between syncs>,
                            min_peers = <minimum peers>,
                            val_data = <validation dataset>,
                            val_batch_size = <validation batch size>,
                            node_weightage = <relative weightage of node’s model weights>)
~~~

sync_interval specifies the synchronization interval,

min_peers specifies the minimum number of nodes for model synchronization,

val_data specifies the validation data set,

val_batch_size specifies the validation batch size,

model_name specifies the name of the model,

node_weightage specifies the relative weightage to be given to model weights of this node

##### Parameter tuning

For some of the scenarios we tuned model hyperparameters. For some scenarios we also tuned Swarm Learning parameters to get better performance, for example higher sensitivity.

For AML **Fig. 2e**, **Extended Data Fig. 2** and **Fig. 2f**, dropout rate is reduced to 10% to get better performance. For AML **Fig. 2b**, **Extended Data Fig. 1**, dropout rate is reduced to 10% and increased the Epochs to 300 to get better performance. We also used the adaptive_rv parameter in the Swarm Learning API to adjust the merge frequency dynamically based on model convergence to improve the training time. For TB and COVID-19 tests dropout rate is reduced to 10% for all scenarios. For the TB scenarios in **Extended Data Fig. 7a,b**, the node_weightage parameter of Swarm Learning callback API is used to give more weightage to the nodes that have higher case samples.

##### Infrastructure layer

###### Description of the hardware architecture applied for simulations

For all simulations provided in this project we used 2 HPE Apollo 6500 Gen 10 server, each with 4 Intel(R) Xeon(R) CPU E5-2698 v4 @ 2.20GHz, a 3.2 TB hard disk drive, 256 GB RAM, 8 Tesla P100 GPUs, 1GB network interface card for LAN access and infiniBand FDR for high speed interconnect and networked storage access. The Swarm Network is created with 3 nodes, each node is a docker container with 1 GPU. Multiple experiments were run in parallel using the above described configuration.

Overall, we performed 6,139 analyses including six scenarios for all three AML datasets, nine scenarios for Tb and 10 scenarios for COVID-19. We performed 5 to 100 permutations per scenario, each permutation took approximately 30 minutes, which resulted in a total of 3069,5 compute hours.

###### The Swarm learning framework, library, distributed ML and blockchain technologies

Swarm Learning builds on top of two proven technologies — distributed ML and blockchain. Distributed ML is leveraged to train a common model across multiple nodes with a subset of the data located at each node — commonly known as the data parallel paradigm in ML — though without a central parameter server. Blockchain lends the decentralized control, scalability, and fault-tolerance aspects to the Swarm Network system to enable the framework to work beyond the confines of a single enterprise.

The Swarm Learning library is a framework to enable decentralized training of ML models without sharing the data. The Swarm Learning framework is designed to make it possible for a set of nodes — each node possessing some training data locally — to train a common ML model collaboratively without sharing the training data itself. This can be achieved by individual nodes sharing parameters (weights) derived from training the model on the local data. This allows nodes to maintain the privacy of their raw data. Importantly, in contrast to many existing federated learning models, a central parameter server is omitted in Swarm Learning.

The nodes that participate in Swarm Learning, register themselves with the Swarm Network implicitly using the callback API. Here, the Swarm Network interacts with other peers using blockchain for sharing parameters and for controlling the training process. On each node, a simple Swarm callback API has to be used to enable the ML model with Swarm Learning capacities (see also code presented below). The Swarm container has to be configured to interact with the Swarm Network (network i/p and port configuration). All other complexities of setting up network, registration, parameter sharing, and parameter merging are taken care of by the Swarm callback API and the Swarm Network infrastructure.

Parameters shared from all the nodes are merged to obtain a global model. Moreover, the merge process is not done by a static central coordinator or parameter server, but rather a temporary leader chosen dynamically among the nodes is used to perform the merge, thereby making the Swarm network decentralized. This provides a far greater fault-tolerance than traditional centralized-parameter-server-based frameworks. All the nodes can perform the role of training and merging, thereby maximising the usage of local compute. The Swarm Network implicitly controls this.

The HPE Swarm Learning library contains 2 containers, the Swarm Network container and the Swarm ML container.

The Swarm Network container includes 1) software to setup and initialize the Swarm Network, 2) management commands to control the Swarm Network, and 3) start/stop Swarm Learning tasks. This container also encapsulates the blockchain software.

The Swarm ML container includes software to support 1) decentralized training, 2) integration with ML frameworks, and 3) it exposes APIs for ML models to interact with Swarm Learning.

For any ML model to be applied to Swarm Learning, it needs to be modified using the Swarm callback API. The callback API provides options to control the Swarm Learning processes. To convert a ML program into a Swarm ML program the following steps have to be performed:

1. Import the SwarmCallback class from the swarm library

~~~
from swarm ‘import SwarmCallback’
~~~

SwarmCallback is a custom callback class that is built on the Keras Callback class.

2. Instantiate an object of the SwarmCallback class:

~~~
swarm_callback = SwarmCallback(min_peers = <peer count>,
                                sync_interval = <interval>,
                                use_adaptive_sync = <bool>,
                                val_batch_size = <batch size>,
                                val_data = <either a (x_val, y_val) tuple or a generator>
                                node_weightage = <relative weightage of node’s model weights>).
~~~

In this context, ‘min_peers’ specifies the minimum number of network peers required to synchronize the insights, ‘sync_interval’ specifies the number of batches after which a synchronization is performed, ‘use_adaptive_sync’ specifies whether the *adaptive sync interval* feature should be used for tuning the sync interval. This feature is turned off by default; ‘val_batch_size’ specifies the size of each validation batch; ‘val_data’ specifies the validation dataset. It can be either a (x_val, y_val) tuple or a generator;

3. Pass the object to the list of callbacks in Keras training code: model.fit(…, callbacks = [swarm_callback]). SwarmCallback can be included along with other callbacks also:

~~~
es_callback = EarlyStopping(…);
model.fit(…, callbacks = [es_callback, swarm_callback])
~~~

###### The Swarm Learning architecture principles

The Swarm Learning framework has two major components, 1) the Swarm ML component runs a user-defined Machine Learning algorithm, and 2) the Swarm Network component forms the Swarm Network based on a blockchain network.

The Swarm ML component is implemented as an API available for multiple popular frameworks such as TensorFlow, Keras, Pytorch. This API provides an interface that is similar to the training APIs in the native frameworks familiar to data scientists. Calling this API automatically inserts the required hooks for Swarm Learning so that nodes seamlessly exchange parameters and subsequently continue the training after setting the local models to the globally merged parameters. With a few simple code changes, the entire network learns as one cohort, with all the complexities of control and data flow taking place in an automated fashion.

Within the Swarm Network component each Swarm ML component interacts with each other using the Swarm Network component’s blockchain platform to maintain global state information about the model that is being trained and to track the training progress. The Swarm Network components use this state and progress information to coordinate the working of the Swarm learning. The Swarm Network is responsible for keeping the decentralized Swarm network in a globally consistent state. The Swarm Network ensures that all operations and the corresponding state transitions are performed in a synchronous manner. Both, state and supported operations of the system are encapsulated in a blockchain smart contract. The Swarm Network contains the logic to elect the leader of the Swarm for every synchronization, implement fault-tolerance, and self-healing mechanisms, along with signaling among nodes for commencement and completion of various phases.

The Swarm Learning framework is designed to run on both commodity and high-end machines, supporting a heterogeneous set of infrastructure in the network. It can be deployed within and across data centers.

In contrast to federated learning with star topology and a centralized coordinator, Swarm Learning can support multiple topologies including fully connected, mesh, star, tree and hybrid topologies. This flexibility provides multiple options to cater into different use cases.

###### The Swarm Learning process

Swarm Learning provides a callback API to enable swift integration with multiple frameworks. This API is incorporated into the existing ML code to quickly transform a stand-alone ML node into a Swarm Learning participant in a non-intrusive way. It offers a set of commands (APIs) to manage the Swarm Network and control the training.

The Swarm learning process is as follows:

The Swarm Learning process begins with enrollment of nodes with Swarm Network, which is done implicitly by Swarm callback function when the callback is constructed. During this process, the relevant attributes of the node are stored in the blockchain ledger. This is a one-time process.

Nodes will train the local copy of the model iteratively using private data over multiple epochs. During each epoch, the node trains its local model using one or more data batches for a fixed number of iterations. It regularly shares its learnings with the other Swarm nodes and incorporates their insights. Users can control the periodicity of this sharing by defining a Synchronization Interval in Swarm callback API. This interval specifies the number of training batches after which the nodes will share their learnings.

At the end of every synchronization interval, when it is time to share the learnings from the individual models, one of the Swarm nodes is elected as a “leader” using the leader election logic. This leader node collects the model parameters from each peer node and merges them. The framework supports multiple merge algorithms such as mean, weighted mean, median, and so on. Each node then uses these merged parameters to calculate various validation metrics. These results are compared against the stopping criterion and if it is found to be met, the Swarm Learning process is halted. Else the nodes use the merged parameters to start the next training batch.

Swarm Learning library uses blockchain smart contracts to define the leader election logic and the merge algorithm. The blockchain smart contracts prevents attacks from semi-honest or dishonest participants.

##### Quantification and Statistical Analysis

We evaluated binary classification model performance with sensitivity, specificity, accuracy and f1-score metrics. Sensitivity, specificity, accuracy and f1-score were determined for every test run. The 95% confidence intervals of all performance metrices were estimated using the boostrapping approach^74^. For AML and ALL, 100 permutations per scenario were run for each scenario. For TB the performance metrics were collected by running 10 permutations for scenarios 1 to 4 and 5 permutations for scenarios 5 to 10. For COVID-19 the performance metrics were collected by running 20 permutations for each scenario. All metrics are listed in **Supplementary Tables 3 and 4.**

Differences in performance metrics were tested using the Wilcoxon signed rank test with continuity correction (Individual Comparisons by Ranking Methods, Frank Wilcoxon, https://sci2s.ugr.es/keel/pdf/algorithm/articulo/wilcoxon1945.pdf). All test results are provided in **Supplementary Table 5**.

To run the experiments, we used Python version 3.6.9 with Keras version 2.3.1 and Tensorflow version 2.2.0-rc2. We used scikit-learn library version 0.23.1^75^ to calculate values for the metrics. Summary statistics and hypothesis tests were calculated using R version 3.5.2 (R: A language and environment for statistical computing, http://www.R-project.org/., 2015). Calculation of each metric was done as follows:

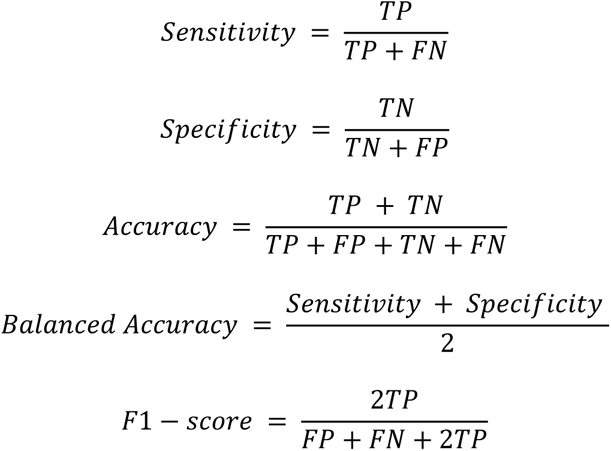

where TP=True Positive, FP=False Positive, TN=True Negative, FN=False Negative

##### Data visualization

The classification report and confusion matrix was generated with scikit-learn APIs for each permutation. Measurements of sensitivity, specificity and accuracy of each permutation run was read into a table in Excel using Power Query and used for visualization for the different scenarios in Power BI [Version: 2.81.5831.821 64-bit (Mai 2020)] with Box and Whisker chart by MAQ Software (https://appsource.microsoft.com/en-us/product/power-bi-visuals/WA104381351).

### Data and software availability

Processed data can be accessed via the SuperSeries GSE122517 or via the individual SubSeries GSE122505 (dataset A1), GSE122511 (dataset A2) and GSE122515 (dataset A3). Dataset B consists of the following series which can be accessed at GEO: GSE101705, GSE107104, GSE112087, GSE128078, GSE66573, GSE79362, GSE84076, and GSE89403. Furthermore, it contains the Rhineland study. This dataset is not publicly available because of data protection regulations. Access to data can be provided to scientists in accordance with the Rhineland Study’s Data Use and Access Policy. Requests for further information or to access the Rhineland Study’s dataset should be directed to. Dataset C contains dataset B and additional samples for COVID-19. These datasets are made available at the European Genome-Phenome Archive (EGA) under accession number EGAS00001004502, which is hosted by the EBI and the CRG.

The code for preprocessing and for predictions can be found at GitHub (https://github.com/schultzelab/swarm_learning).

### Supplementary Tables

Supplementary Table 1: Overview over all sample numbers and scenarios

Supplementary Table 2: Dataset annotations of Dataset A, B and C

Supplementary Table 3: Prediction results for all scenarios and permutations

Supplementary Table 4: Summary statistics on all prediction scenarios

Supplementary Table 5: Statistical tests comparing single node vs. swarm predictions

Supplementary Table 6: Covid 19 Patient characteristics

